# Capture, mutual inhibition and release mechanism for aPKC-Par6 and its multi-site polarity substrate Lgl

**DOI:** 10.1101/2024.09.26.615224

**Authors:** Christopher P. Earl, Mathias Cobbaut, André Barros-Carvalho, Marina E. Ivanova, David C. Briggs, Eurico Morais-de-Sá, Peter J. Parker, Neil Q. McDonald

## Abstract

The mutually antagonistic kinase-substrate relationship between the apical aPKC-Par6 heterodimer and the basolateral substrate Lgl is key to the establishment and maintenance of cell polarity across metazoa. Although aPKC-Par6 can phosphorylate Lgl at three serine sites to exclude it from the apical domain, paradoxically, aPKC-Par6 and Lgl can also form a stable kinase-substrate complex whose function remains unclear and with conflicting roles proposed for Par6. We report the structure of human aPKCι-Par6α bound to full-length Llgl1, captured through an aPKCι docking site and a Par6^PDZ^ contact. This soluble tripartite complex traps a phospho-S663 Llgl1 intermediate bridging between aPKC and Par6, impeding phosphorylation progression. Thus, aPKCι is effectively inhibited by Llgl1^pS663^ whilst Llgl1 is captured by aPKCι-Par6. Mutational disruption of Lgl-aPKC interaction impedes complex assembly and Lgl phosphorylation, whereas disrupting the Lgl-Par6^PDZ^ contact promotes complex dissociation and completion of Lgl phosphorylation cycle. We incorporate these findings into a Par6^PDZ^-regulated substrate capture-and-release model that we demonstrate requires binding by Cdc42-GTP and the apical partner Crumbs to drive complex disassembly. Our results provide an explanation for the opposing roles of Par6 underpinning the spatial control of aPKC-Par6 activity by Lgl relevant to polarised membrane contexts across multi-cellular organisms.

## Introduction

Apical-basal polarity in epithelial cells is formed by the action of a conserved network of partitioning defective (Par) proteins and their multi-protein complexes ^1–3^. This network exhibits properties of mutual membrane exclusion (also known as mutual antagonism) and feedback to form polarised membrane domains with unique identities and sizes ^4^. How these emergent and dynamic properties arise from the formation of multi-protein Par assemblies is not fully understood, but phosphorylation of membrane-bound substrates is believed to be key. Within the apical membrane domain, the kinase-substrate relationships of the atypical protein kinase C (aPKC in *Drosophila* and two isoforms aPKCι/aPKCζ in human) dominate ^5–9^. aPKC associates with the PDZ-protein Par6 to phosphorylate substrates such as Par1, Par2 and Par3, that frequently bear an F-X-R docking motif, driving them off apical membranes ^10^. The precise contribution of Par6 in this process is unclear however, with both activating and inhibitory roles towards aPKC kinase activity having been proposed ^11,12^.

Polarity components oppose and repress aPKC activity in the cytoplasm or at lateral membranes restricting aPKC-Par6 activation to apical membranes ^1,11^. A well characterised example of a spatially-controlled antagonist of aPKC is Lethal (2) giant larvae (Lgl in *Drosophila* and Llgl1/2 in mammals; in this study we will use Lgl generically across all species to refer to Lgl/Llgl1/Llgl2, other than where we explicitly refer to Lgl in experiments in *Drosophila*). Lgl restrains aPKC activity except at the apical membrane where Lgl is removed directly by aPKC-Par6 through phosphorylation ^1,11^. Lgl has a double ß-propeller structure and contains multiple phosphorylation sites for aPKC-Par6 with at least three serine phospho-acceptor sites (S655, S659, S663 in human Llgl1, within a segment defined hereafter as the P-site) that map within a key membrane-binding loop ^13^. These highly conserved sites are required for efficient cortical displacement of Lgl but are both functionally and kinetically distinct ^14–17^. Active apical aPKC suppresses apical action of Lgl by phosphorylating its P-site, and displacing it from the membrane ^18,19^. Lgl is consequently localised to the basolateral membrane at steady state (together with Scribble and Discs Large (Dlg)) in a manner mutually exclusive with aPKC localisation ^18,20^. This reciprocal localisation depends on Lgl phosphorylation by aPKC as an Lgl mutant lacking the three phospho-sites invades the apical membrane ^14,20^. Paradoxically, Lgl forms a stable tripartite complex with aPKC-Par6 in a manner mutually exclusive with Par3, suggesting a more complex regulation than a simple hit-and-run phosphorylation mechanism ^9,21^. The molecular mechanism for mutual inhibition is not understood despite its key role in many cell polarity contexts including epithelial polarity, asymmetric cell division, and neuronal polarisation.

To understand the antagonism between Lgl substrate and aPKC-Par6 kinase, the basis for multi-site phosphorylation and how this stable three-way kinase-substrate complex is assembled, we determined a cryo-EM structure of an aPKC-Par6-Lgl tripartite complex. The structure reveals the intricate interplay between aPKC-Par6 and its substrate Lgl, which together with *in vitro* and *in vivo* data support a capture and release mechanism involving a stable phospho-intermediate. In this mechanism, the Par6^PDZ^ domain is uncovered as the missing molecular link explaining exquisite substrate targeting of Lgl, the mutual inhibition of aPKC and its role as an apical sensor coupled to an allosteric release mechanism. Thus, we provide a description of a near complete kinase-substrate multi-site phosphorylation reaction cycle.

## Results

### Structure of a mutually-antagonised stalled polarity complex

To obtain the structure of an aPKC-Par6-Lgl tripartite complex, we expressed and purified a complex containing human aPKCι, Par6α, and Llgl1, from Freestyle HEK293-F cells (Extended data Figure 1a). Formation of a soluble complex was confirmed by size-exclusion chromatography. The structure of human aPKCι-Par6α-Llgl1 polarity complex was then determined at a nominal resolution of 3.44Å using cryo-EM (Extended data Figure 1b-d). Maps were of sufficient quality to reliably dock the Par6 PDZ domain (salmon, PDB code 1RZX), the aPKCι kinase domain (yellow, PDB code 5LI9) and a crystal structure of full-length Llgl2 (blue, PDB code 6N8Q) with the readily recognisable double beta-propeller (Figure 1a-b, Extended data Figure 1e-f) ^5,13,22^. After fitting each component and adjusting their sequences to the correct isoform, the molecular interfaces were rebuilt *de novo*, including all conserved parts of the previously unseen Llgl1 membrane-binding loop (10-11) harbouring the P-site (Figure 1c-e). This gave a reliable atomic model for the aPKCι-Par6α-Llgl1 tripartite assembly showing reciprocal interactions between all three components (Figure 1b). The nucleotide pocket was occupied with AMP-PNP added to the sample prior to grid preparation to stabilize the kinase core (Figure 1b, Extended data Figure 1g) and two known phospho-threonine sites in aPKC (pT412^aPKCι^ and pT564^aPKCι^) and one phospho-serine in Llgl1 (Llgl1^pS663^) were unequivocally identified (Extended data Figure 1h-k). Regions towards the periphery of the cryo-EM map were less well resolved, including the pseudo-substrate (PS) membrane-binding element of aPKCι, as well as the PB1 domains of aPKCι and Par6α (Extended data Figure 1l). Additional density proximal to the αC-helix of the kinase domain N-lobe was observed in cryo-EM 2D class averages and 3D reconstructions (Extended data Figure 1l), which matched the overall shape and size of the aPKCι C1 domain but could not be fit reliably.

**Figure 1:**
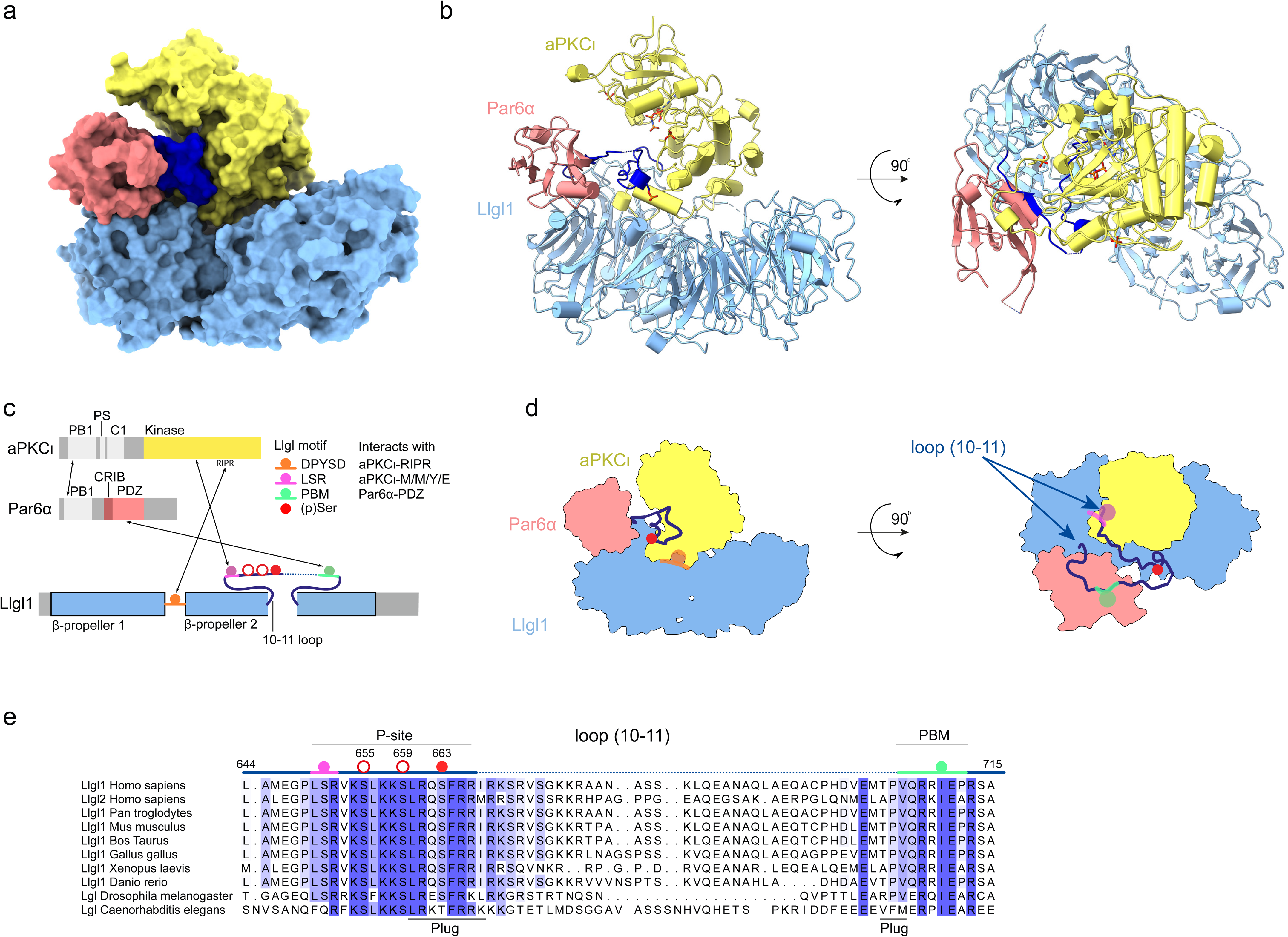
Cryo-EM structure of an antagonised aPKCι-Par6α-Llgl1 polarity complex. (a) Surface rendering of the aPKCι-Par6α-Llgl1 cryo-EM structure. Surfaces for individual components are coloured differently, Par6 (flesh), aPKCι (yellow) and Llgl1 (light blue) and the Llgl1 (10-11) loop containing the P-site (navy) (b) Ribbon diagram of the aPKCι-Par6α-Llgl1 complex with Par6α (flesh), aPKCι (yellow) and Llgl1 (light blue) and the Llgl1 (10-11) loop containing the P-site (navy). A stick representation for AMP-PNP is shown indicating the aPKCι nucleotide-binding site as well as the three phospho-residues aPKCι ^pT412,^ aPKCι ^pT564^ and Llgl1^pS663^ (c) Schematic of key interactions mapped onto the domain structures for each component. Greyed out segments indicate regions not defined in the final model (d) Schematized aPKCι-Par6α-Llgl1 structure showing two orthogonal slices through the structure with similar view to (b) mapping the approximate location of crucial interaction contacts and phospho-acceptor serine residues (e) Alignment of loop (10-11) sequences for Lgl homologues indicating conservation at opposing ends of the membrane-binding loop. Phospho-acceptor sites and interaction contacts are coloured as per panel (c).

The resulting aPKCι-Par6α-Llgl1 structure reveals the basis for coordinated capture of Llgl1 by Par6 and aPKCι, leading to a stably associated (stalled) phospho-intermediate of Llgl1. Multiple interactions stabilise the complex, mapping onto the second Llgl1 ß-propeller and its central membrane-binding loop (10-11) that contains the P-site (Figure 1c-e). The second ß-propeller of Llgl1 makes the largest contact with the aPKCι kinase domain (aPKCι^KD^) insert (residues G457 to D468). This insert is located between the F and G helices of the kinase C-lobe extending away from the kinase to penetrate into the ß-propellor cavity (Figure 2a). This contact involves both acidic and hydrophobic contacts to bury a total surface area of ∼2374Å^2^ (calculated at https://www.ebi.ac.uk/pdbe/pisa/) providing an extensive landing pad for the aPKCι^KD^. A second Llgl1-aPKCι^KD^ contact involves the aPKC RIPR motif (residues R480 to R483) in the loop between the αG and αH-helix also at the base of the kinase C-lobe. This element straddles a complementarily charged DPYSD motif on Llgl1 located within loop (8-9) (Figure 2b). This satisfyingly explains our previous observations that the RIPR motif is a necessary contact site for Llgl1/2 phosphorylation ^23^. Both these contacts help orient the aPKCι kinase domain with its substrate-binding cleft facing towards and anchoring part of the ∼72 aa-long membrane-binding loop (10-11) of Llgl1 containing the P-site. Residues L640-S641-R642 at the amino-terminus of the Llgl1 P-site engage a high affinity docking site on aPKCι that was previously shown to be occupied by Par3 FXR-motif (Figure 2a) ^5^. The C-terminus of Llgl1 loop (10-11) is bound to the Par6 PDZ domain (Par6α^PDZ^) through a previously unrecognised internal **P**DZ-**<u>b</u>**inding **<u>m</u>**otif (Llgl1^PBM^) spanning residues V706-P712 (Figure 1e). The Llgl1^PBM^ is highly conserved in Lgl homologues explaining the strong sequence constraints at the C-terminal side of Llgl1 loop (10-11). The Llgl1^PBM^ adopts a short ß-strand (Figure 2c) that completes the central Par6 PDZ domain ß-sheet as observed for other PDZ ligands ^22^. Internal PBMs are poorly characterised, but typically consist of a hydrophobic amino acid followed by an acidic residue (mimicking the carboxy-terminal interaction of canonical PBMs) ^24^. The key Llgl1 residue I710 occupies a hydrophobic pocket formed by L169, F171, I173 and M235 within the PDZ cleft and is followed by E711 (Figure 2c). P712 inserts towards the PDZ core disfavouring the PDZ conformation that binds carboxy-terminal PBM motifs with high affinity (see further).

**Figure 2:**
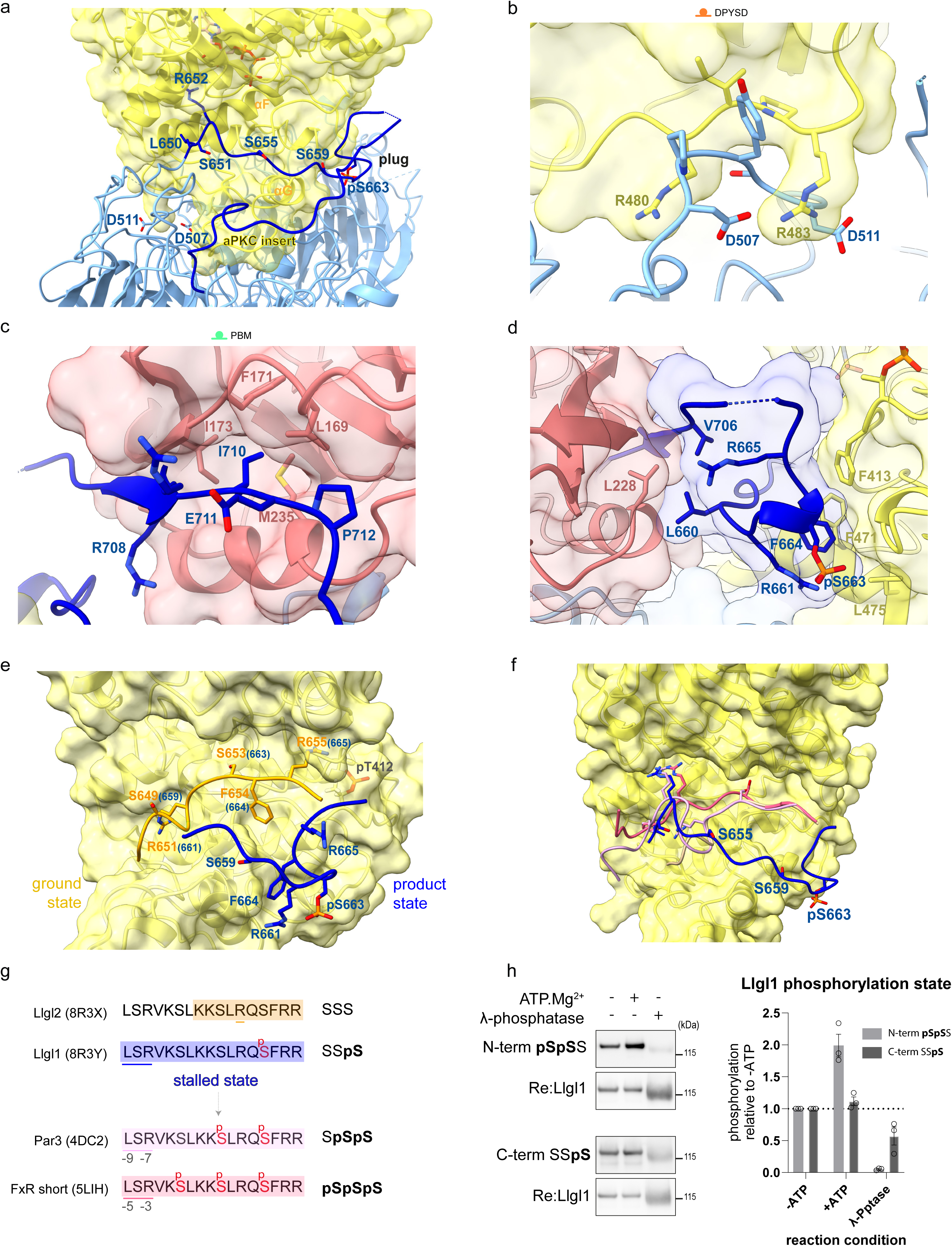
Capture of membrane-binding Llgl1 loop by aPKCι-Par6α controls P-site phosphorylation progression. (a) Extensive contacts between the aPKCι core kinase domain (yellow solid rendering) and the Llgl1 loop (10-11) plug (navy). The location of the P-site phospho-acceptor sites in Llgl1 are indicated (S655, S659 and pS663) together with selected residues from the aPKCι LSR- and RIPR-motifs. The position of the plug domain formed around pS663 is also indicated. (b) Close-up of key residues from the RIPR-motif contact within aPKCι and the reciprocal DPYSD motif in Llgl1. (c) Close-up of key residues from the Par6α PDZ contact with an internal PBM motif within the 10-11 loop of Llgl1. (d) Close up of the molecular “plug” formed by either opposing ends of the Llgl1 loop (10-11) showing key residues close to the LRQSFRRI P-site sequence contributing to a small hydrophobic core or bridging between the aPKCι C-lobe pocket and the Par6α PDZ domain (e) Structural superposition of the Llgl2 P-site peptide crystal structure reported here overlayed with the trajectory of the phosphorylated product peptide from the tripartite cryo-EM structure. The superposition reflects the likely ground state of the Lgl P-site in its unphosphorylated state and the conformational change induced upon phosphorylation of the initial pS663 site referred to as the “stalled” state. (f) Proposed reconstruction of phosphorylation progression trajectory from the unphosphorylated state to the “stalled” state (pS663) to a double (pS659-pS663) then triple phosphorylation state (pS655-pS659-pS663). Previous structures have shown how substrates with an F^-9^XR^-7^ motif drive phospho-acceptor phosphorylation or through an F^-5^XR^-3^ motif (PDB codes 4DC2 and 5LIH). The LSR motif of Llgl1/2 conformation adopts an identical pose to the FSR motif within the Par3 CR3 domain, we propose that release of the “captured” poise (see later) enables rotation of the P-site into the substrate binding pocket for double (pS659-pS663) and subsequent triple phosphorylation (pS655-pS659-pS663). (g) Summary of P-site phosphorylation progression highlighting the stalled state (h) Immunoblot evidence that Llgl1 is sub-stoichiometrically phosphorylated on the N-terminal Ser residues but stoichiometrically phosphorylated on the C-terminal Ser residue within the aPKCι-Par6α-Llgl1 complex. Quantification is shown on the right (n=3 independent *in vitro* assays, represented as mean +/- SEM).

Tethering of either end of the Llgl1 loop (10-11) to aPKCι^KD^ and Par6^PDZ^ respectively as described above has three consequences. First, it separates the extremities of the loop and guides both into a deep cleft formed between aPKCι^KD^ and Par6^PDZ^ (Figure 1a and 2a). Second, it brings two segments of the loop into close proximity to bridge between aPKCι^KD^ and Par6^PDZ^ by forming a molecular “plug” domain nucleated around the single site phosphorylation at Llgl1^pS663^ (Figure 2d). The phospho-serine 663 site (Llgl1^pS663^) identified in the cryo-EM potential map forms a salt bridge with R661^Llgl1^ which packs against the F664^Llgl1^ sidechain. These two residues in turn engage a conserved hydrophobic pocket within the aPKCι^KD^ C-lobe (involving residues F413, F471, L475). The plug domain itself has a small hydrophobic core made up of L660 and V706 and the aliphatic parts of several basic residues (K658/R665/R708) that cross from one side of the cleft to the other.

The third consequence of the tethered ends of Llgl1 loop (10-11) is to trap the P-site in a non-productive conformation for phosphorylation progression. By forming the plug domain around Llgl1^pS663^, the P-site effectively blocks access of S655 and S659 phospho-acceptor sites to the aPKCι substrate cleft and catalytic residues (Figure 2e). The organization of this tripartite structure argues that the aPKCι-Par6α heterodimer captures Llgl1, preventing its critical membrane-interaction motif within the P-site from binding to the plasma membrane. Furthermore, the identification of this inhibited state further argues that additional steps must take place to promote multisite phosphorylation and in turn release Lgl from the complex.

### Evidence for an Lgl multi-site phosphorylation trajectory

The observation that only Llgl1^S663^ in the P-site was phosphorylated within the cryo-EM structure agrees with previous *in vitro* peptide substrate studies that the C-terminal phosphorylation of the P-site is catalytically favoured ^16^ and presented first to aPKC. To support these findings, we investigated how the P-site is presented in its unphosphorylated state. A peptide spanning the Llgl1 P-site binds the aPKCι^KD^ with an apparent affinity (K_d_) of 48nM (Extended data Figure 2a). We then screened for crystals of the Llgl1 and Llgl2 P-site peptides and determined a medium resolution structure for the Llgl2 P-site peptide spanning residues 634-662 bound to aPKCι^KD^ in the absence of nucleotide. Llgl2 P-site is identical to the Llgl1 sequence in this region but is ten residues shorter in numbering. The co-crystal structure of aPKCι^KD^-Llgl2^P-site^ revealed ordered contacts between residues 650-656 of the Llgl P-site, including the C-terminal S653 (equivalent to Llgl1 S663). In this poise, Llgl2^S653^ is bound ready for phospho-transfer, with Llgl2^F654^ in the hydrophobic +1 pocket (Figure 2e and Extended data Figure 2b). The R655 sidechain hydrogen bonds to the G398 carbonyl and a phosphate oxygen from phospho-T412, while R651 sidechain at the -2 position, relative to the phospho-acceptor, lies in a pocket formed between Y419 and E445 of aPKCι (Extended data Figure 2b). The structure confirms the Llgl P-site binds aPKC thereby presenting Llgl2^S653^ (equivalent to Llgl1^S663^) for initial phosphorylation, consistent with the cryo-EM structure. Absent from the X-ray structure is the L640-S641-R642 motif, suggesting that this motif may not be crucial for the initial C-terminal Llgl1/Llgl2 phosphorylation. We propose phosphorylation progression of the two amino-terminal sites can be modelled from previous substrate peptide structures (PDB codes 5LIH and 4DC2) corresponding a short (F^-5^)-X-(R^-3^) motif or a long (F^-9^)-X-(R^-7^) motif binding mode respectively (Figure 2f) ^5^. These modes differ in the register between the bound L-S-R motif and phospho-acceptor sites Llgl1 S655 and S659, which individually fit the (F^-5^)-X-(R^-3^) and (F^-9^)-X-(R^-7^) motif. Superposition of previous substrate structures (PDB codes 4DC2 and 5LIH) indicate the precise conformational movement required to position the two N-terminal phospho-acceptor serine residues S655 and S659 close to the gamma-phosphate of ATP. In each case the phospho-acceptor C-alpha position would need a substantial conformational shift of 9.7Å and 13.4Å respectively from the cryo-EM structure to be available for phospho-transfer. Thus, we can predict the phosphorylation trajectory from snapshots of unphosphorylated peptide (X-ray structure) to a stalled initial site phosphorylation (cryo-EM structure) and predicting the S655 and S659 phosphorylation binding poses based on related substrate peptide structures (Figure 2g).

Although the C-terminal P-site phosphorylation (mono-phosphorylated at Llgl1^S663^) state is the predominant form captured within the complex, further phosphorylation can be driven *in vitro* by adding ATP.Mg^2+^ to the complex. This increases the phosphorylation on the N-terminal S655/659 sites, but not on the C-terminal S663 site as it is already stoichiometrically phosphorylated as judged by immunoblot with antibodies specifically recognizing these phospho-sites (Extended data Figure 2c), supporting our structure of the single mono-phosphorylated state (Figure 2h). Interestingly, previous *in vivo* studies in *Drosophila* have shown that overexpression of the S656A/S660A double mutation (*Drosophila* Lgl numbering for the N-terminal sites) acts as a potent inhibitor of aPKC, but did not clarify the underlying reason for the specific impact of this mutant version ^17,25^. Phosphorylation at the available C-terminal serine would mimic the stalled phospho-intermediate, effectively trapping aPKC-Par6-Lgl in a state equivalent to that observed in our cryo-EM analysis, thereby explaining the increased ability of this mutant to inhibit aPKC.

### Behaviour of Llgl interface mutations *in vitro*

We then explored how different contacts within the complex contribute to stabilization of the stalled enzyme-intermediate state. To do this we prepared disruptive mutations at the different conserved interfaces within Llgl2, a close human homologue to Llgl1 ^26^ whose structure ^13^ and function ^21,27,28^ is better characterised *in cellulo*. We interrogated interaction site mutants of Llgl2 expressed in HEK cells designed from the Llgl1 tripartite structure (Figure 3a) by assessing their impact on the formation of the tripartite complex. Knowledge of the Llgl1 interaction site with the aPKCι RIPR-motif, allowed us to assess the impact of mutations on either partner through engineering charge reversal mutations to disrupt Llgl2 interaction with the aPKCι RIPR-motif. Consistent with our Llgl1-containing complex structure, we observed that formation of a stable complex between aPKCι and Llgl2 required co-expression with Par6α (Figure 3b). When only aPKCι and Llgl2 were co-expressed, very low levels of aPKCι were recovered bound to GFP-Llgl2. Mutating the RIPR motif in aPKCι to DIPD (or a corresponding DPYSD>RPYSR mutation in Llgl2) suppressed the ability of Par6 to stabilize the tripartite complex, validating the structurally observed interface, in line with previous observations ^23^. Evidence that the stability of the three-way complex is governed primarily by the Par6 contact is shown by mutation of the PBM residue in Llgl2 at I700 (hereafter Llgl2^IE>NE^) (Figure 3a). Disrupting this key hydrophobic contact to the Par6 PDZ completely abolished the interaction with aPKCι and Par6 in HEK293 cells (Figure 3c). These data indicate that formation of a stable complex at steady state requires a three-way interaction involving an aPKC kinase domain-Lgl contact (driven by the aPKCι RIPR motif) and crucially a Par6-Lgl contact (driven by the Par6 PDZ domain).

**Figure 3:**
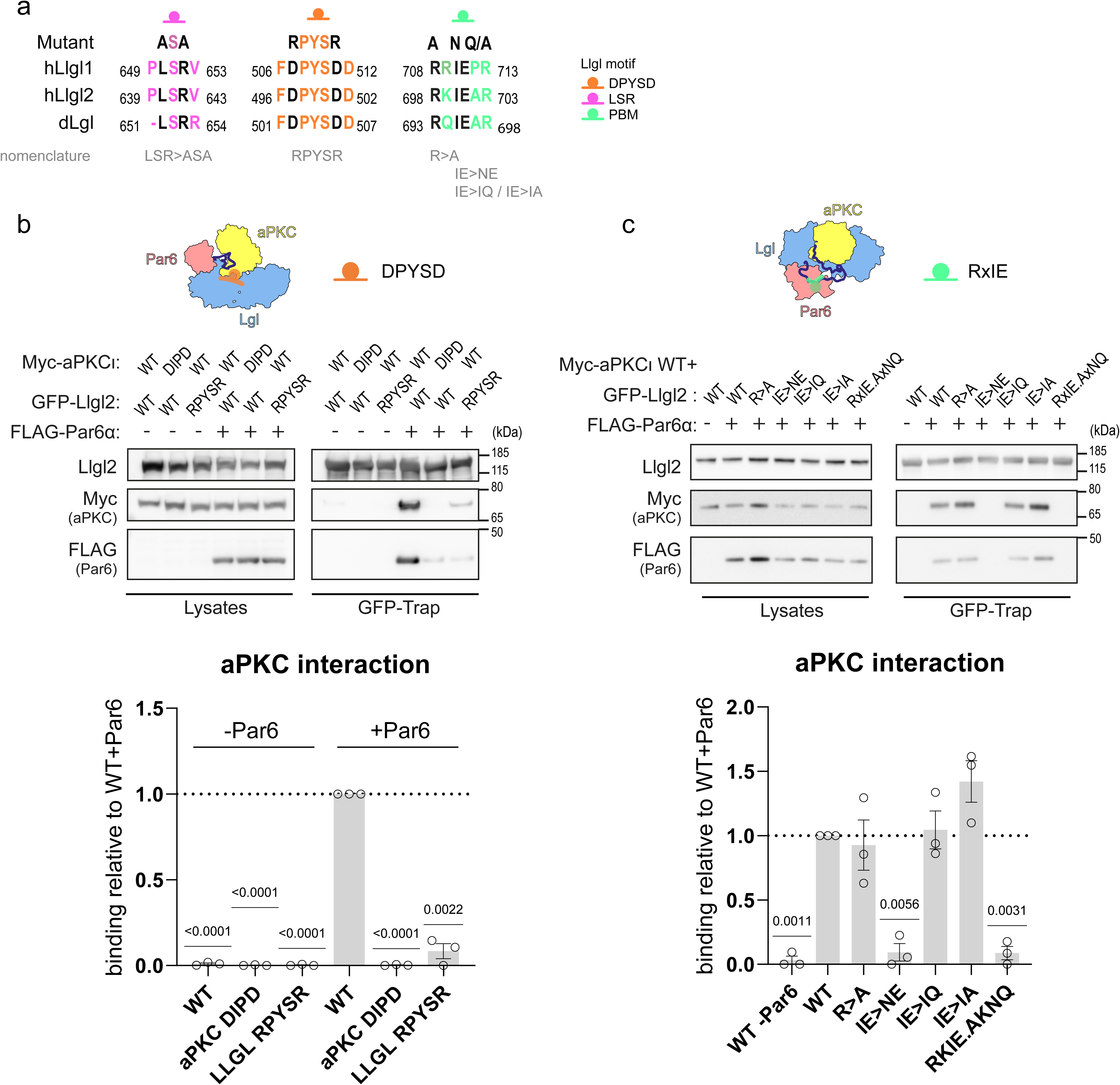
Impact of interface mutations on tripartite complex stability. (a) Lgl docking motifs within the complex and their conservation between isoforms and species. The mutants used to probe the function of each motif are indicated above and their nomenclature used throughout the manuscript shown below in grey. (b) Western blot analysis of GFP-Trap affinity pulldown assays from HEK293 cells expressing different GFP-Llgl2 docking interface mutants, Myc-tagged aPKCι and FLAG-tagged Par6α. Quantification of the western blots is shown directly below the blots (n=3 biological replicates represented as mean +/- SEM, analysed via a two-tailed one-sample t-test) (c). GFP-Trap affinity pulldown assays from HEK293 cells expressing selected GFP-Llgl2 mutants, Myc-tagged aPKCι and FLAG-tagged Par6α. Quantification of the Western blots is shown directly below the blots (n=3 biological replicates represented as mean +/- SEM, analysed via two-tailed one-sample t-test)

### Impact of Lgl mutations in polarised cells and *in vivo*

To explore the behaviour of the validated interface mutants in the context of polarised cells, we expanded on a published phenotype in cultured epithelial cells, whereby wild type Llgl2 overexpression causes a loss of polarity ^21^. We engineered human DLD1 epithelial colorectal cancer cells to overexpress either wild type (WT) GFP-Llgl2 or variants harbouring interface mutations that we had characterised in the kinase-docking DPYSD motif (Llgl2^DPYSD>RPYSR^), the PBM motif (Llgl2^IE>NE^) as well as the substrate docking LSR motif (Llgl2^LSR>ASA^). Each mutant expression was driven from a doxycycline-inducible promotor. In the absence of doxycycline, these cells display a largely polarized phenotype with intact ZO-1 staining indicating properly formed tight junctions and polarised cell contacts (Figure 4a,c). Inducing WT Llgl2 expression results in a dominant membrane localization of the overexpressed protein and loss of polarity, as evidenced by a complete loss of intact ZO-1 staining (Figure 4a,c). By contrast, when expression of the PBM Llgl2^IE>NE^ mutant is induced, no dominant phenotype is observed, and levels of intact ZO-1 staining are similar to the non-induced condition (Figure 4b,c). Although a subfraction of the mutated protein does localize to the plasma membrane, the majority remains in the cytoplasm and notably also clusters in foci that stain positive for the trans-Golgi marker TGN46 (Extended data Figure 3a). Lgl has initially been shown to localize at the Golgi, and the localization of this mutant may therefore reflect a distinct state of this protein ^19^. By contrast, overexpressing either the kinase domain contacting Llgl2^DPYSD>RPYSR^ or Llgl2^LSR>ASA^ variants – or a non-phosphorylatable triple serine to alanine mutant (Llgl2^SSS>AAA^) – resulted in a similar phenotype to that seen with overexpressed WT Llgl2, with a predominant membrane localization and loss of intact ZO1 staining (Extended data Figure 3b,c).

**Figure 4:**
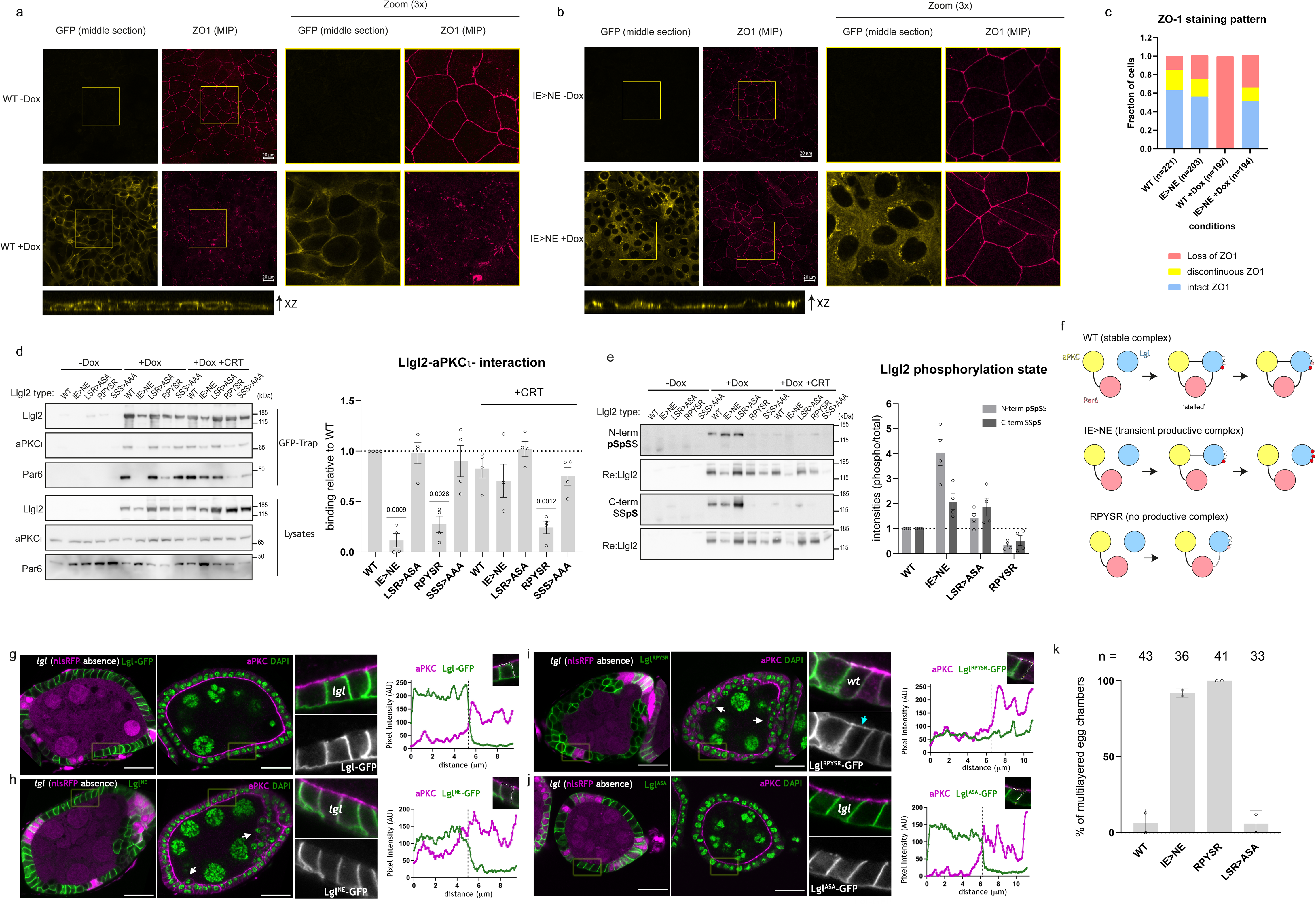
*In cellulo* and *in vivo* characterisation of aPKC-Par6-Lgl complex interface mutants. (a) Localization of WT or (b) Llgl2^IE>NE^ in DLD1 cells and the effect on ectopic protein expression on DLD1 epithelial organization. WT or Llgl2^IE>NE^ were expressed in DLD1 cells via doxycycline (Dox) induction. Llgl2 and ZO-1 localization were followed using confocal microscopy. Representative micrographs shown of one of three independent biological replicates (c) Quantification of cells in (a,b) with intact ZO-1, discontinuous ZO-1 or loss of ZO-1. Representative experiment of one of three biological replicates (d) Complex formation between Llgl2 and aPKCι-Par6. Cells expressing WT or mutant forms of GFP-tagged Llgl2 were lysed with or without pre-treatment with the aPKCι inhibitor CRT0329868 and complex formation between Llgl2 and aPKCι-Par6 was followed by GFP-trap (quantification on the right n=4 biological replicates represented as mean +/- SEM, analysed via a two-tailed one-sample t-test). The GFP-trap Par6 signal was collected on a separate membrane to improve detectability. (e) Phosphorylation state of ectopically expressed Llgl2 in DLD1 cells. Cells expressing WT or mutant forms of GFP-tagged Llgl2 were lysed with or without pre-treatment with the aPKCι inhibitor CRT0329868 and phosphorylation of aPKCι target sites was followed using two antibodies, recognizing the two N-terminal and C-terminal phosphorylation sites of Llgl2 respectively (quantification on the right n=4 biological replicates represented as mean +/- SEM). (f) Proposed effects of mutations in Llgl2 on its phosphorylation and complex formation with aPKCι (for details see text). (g-j) Confocal microscopy images of mosaic egg chambers of *lgl* mutant follicle cell clones (absence of nlsRFP, left panel), expressing the indicated Lgl versions in the *Drosophila* follicular epithelium and stained for DAPI (green, central panel) and aPKC (magenta, central panel). Close-ups and a plot of cortical pixel intensity from the basal side of the denoted region in follicle cells are shown on the right. (k) Graph shows frequency (mean ± SD of epithelial multilayering in egg chambers with mutant clones (larger than 1/4 of the egg chambers) expressing the indicated Lgl versions. n is number of egg chambers from 2 independent experiments.

In parallel with these observations, we studied tripartite complex formation and Llgl2 phosphorylation at steady state in these cells. The WT Llgl2 protein forms a stable complex with aPKCι and Par6, similar to a non-phosphorylatable Llgl2^SSS>AAA^ mutant. This indicates that overexpressed Llgl2 accumulates endogenous aPKCι-Par6 into a stalled complex (Figure 4d). The Llgl2^IE>NE^ mutant on the other hand did not form a stable complex with aPKCι-Par6 at steady state in the DLD1 cells (Figure 4d), consistent with our observations in HEK293 cells. However, compared to the WT protein, the PBM mutant Llgl2^IE>NE^ shows increased phosphorylation, predominantly at the N-terminal serine sites (Figure 4e). This observation argues that the Llgl2^IE>NE^ protein forms only a transient phosphorylation complex with aPKCι, and conversely that stable interaction of WT Llgl2 with Par6 suppresses catalytic activity – mainly at the N-terminal Ser-645/649 phosphorylation sites. Additionally, the transient interaction of the Llgl2^IE>NE^ mutant allows for the phosphorylation of endogenous Llgl1/2 by aPKC-Par6, while the overexpressed WT mutant supresses this (Extended data Figure 3d). The third mutant Llgl2^LSR>ASA^ displays slightly elevated levels of phosphorylation skewed towards its C-terminal phosphorylation sites, while exhibiting wild type levels of N-terminal Ser phosphorylation and tripartite complex formation (Figure 4d,e). The fact that this mutation results in higher relative levels of C-terminally phosphorylated Llgl2 argues that there is an increased turnover rate compared to the WT protein and supports the notion that the LSR motif aids phosphorylation of the N-terminal Ser-645/649 sites, but appears dispensable under these conditions. The Llgl2^DPYSD>RPYSR^ mutant breaks the aPKCι docking contact, leading to reduced Llgl2 binding to aPKCι and reduced levels of phosphorylation (Figure 4d,e). In contrast to the HEK293 cells in the context of co-expression with Par6α, in DLD1 polarised cells the loss of the aPKC-Llgl2 contact is less disruptive to the three-way complex, with residual binding likely mediated by Par6. This is supported by the observation that treatment with an aPKC-selective chemical inhibitor CRT0329868 ^29^ traps the PBM-defective Llgl2^IE>NE^ variant in the tripartite complex, whereas it has no such effect on the kinase contact Llgl2^DPYSD>RPYSR^ variant (Figure 4d), highlighting that binding to the kinase domain is impaired in the latter case and not in the former.

Taken together, these data suggest that overexpressed WT Llgl2 forms stable complexes with aPKCι-Par6, trapping it in an auto-inhibited complex with low turnover (Figure 4f, top row). The PBM mutant Llgl2^IE>NE^ instead forms a transient complex with aPKCι that is efficiently phosphorylated and released, resulting in a higher stoichiometry of phosphorylation and cytosolic protein that accumulates in the Golgi compartment (Figure 4f, middle row). The kinase docking Llgl2^DPYSD>RPYSR^ variant, in contrast, displays reduced levels of complex formation in the DLD1 cells and reduced capacity to form a productive enzyme-substrate complex with aPKCι-Par-6, resulting in reduced phosphorylation (Figure 4f, bottom row). Produced in excess, this mutant can disrupt polarity likely because it shows residual binding to aPKC, and cannot be cleared from membranes by phosphorylation, resulting in an increased pool of membrane-bound Llgl-2 acting on aPKC and other downstream effectors such as Myosin II ^30^.

To characterise these Lgl mutants *in vivo,* we investigated whether these Lgl mutants could support epithelial apical-basal polarity in the monolayered follicular epithelium of the *Drosophila* ovary. This is a well-defined system to probe the *in vivo* role of apical-basal polarity proteins such as Lgl, for which loss of function alleles induce the formation of multi-layered epithelium ^31,32^. We performed rescue experiments in mosaic tissue containing *lgl* null follicle cell clones to analyse the ability of the aforementioned interface mutants to sustain apical-basal organization (mutants defined in Figure 3a). The mutated Lgl variants were tagged with a C-terminal GFP tag and expression was driven specifically in the follicular epithelium using the UAS/GAL4 system ^33^. We then imaged fixed mosaic *Drosophila* egg chambers co-stained for aPKC where the localization of each Lgl variant was detected by the GFP-signal and where *lgl* null cells were identified by absence of nlsRFP (RFP with a nuclear localisation sequence). Expressing the GFP-tagged versions of WT Lgl or the Lgl^LSR>ASA^ mutant restored monolayered architecture in the absence of endogenous Lgl (Figure 4g,j,k), whereas the kinase docking mutant Lgl^DPYSD>RPYSR^ and the PBM mutant Lgl^IE>NE^ showed a large frequency of egg chambers with the multi-layered phenotype (Figure 4h,i,k).

The finding that the Lgl^DPYSD>RPYSR^ and the Lgl^IE>NE^ mutants are unable to support intact tissue polarity indicates the importance of regulated aPKC-Par6-Lgl complex assembly and disassembly to support apical-basal polarity. However, these mutants have distinct functional and localization properties. First, the PBM mutant Lgl^IE>NE^ retains some ability to sustain apical-basal polarization as indicated by the polarized enrichment of aPKC in monolayered patches that lack endogenous Lgl but express Lgl^IE>NE^ (Figure 4h, closeup). In contrast, the mutant disrupting the kinase domain contacts, Lgl^DPYSD>RPYSR^, is unable to support polarity. Second, whereas Lgl^IE>NE^ is cleared from the apical compartment and restricted to the basolateral cortex in polarized cells (Figure 4i panel C), the Lgl^DPYSD>RPYSR^ mutant distributes all over the cell, invading the apical domain even in control nlsRFP positive cells (Figure 4j, closeup). Apical invasion by the Lgl^DPYSD>RPYSR^ mutant is consistent with the lack of Lgl phosphorylation, since it cannot form a productive tripartite complex with aPKC-Par6 promoting its removal. Taken together, these data stress the *in vivo* significance of the stability of the aPKC-Par6-Lgl complex and the dynamics of complex turnover in relation to the phenotype. Inappropriate early release of the Par6 PDZ domain in the Lgl^IE>NE^ mutant likely promotes untimely complex disassembly and reduces the ability of Lgl to antagonize aPKC to maintain apical-basal organization.

### Cdc42 and Crb trigger ATP exchange and complex disassembly

Having established that the Par6^PDZ^ engagement is crucial for stalling the complex, we investigated possible triggers for its release. A prime candidate is Cdc42-GTP that has been shown to induce a conformational switch in Par6^PDZ^ to alter its PDZ specificity for C-terminal PBM partners ^22,34,35^ (Figure 5a). However, the biological significance and context for this allosteric switch mechanism has not been demonstrated. Comparison of the PDZ domain conformation bound to the internal Llgl1 PBM shows close similarity to the previously reported complex with an internal Pals1 PBM ligand ^22^ (Figure 5a). However, comparison to the Crumbs3 (Crb3) C-terminal PBM bound complex revealed a steric clash between P712 of Llgl1 (within the PBM at I710-E711) and a lysine in Par6 (K165 in *Drosophila* Par6, equivalent to K162 of human Par6). This lysine is part of an allosteric dipeptide switch motif (L164/K165 in *Drosophila* Par6 and H161/K162 in human Par6) that has been proposed to bind Crb3 with high affinity in the presence of Cdc42-GTP ^35^ (Extended data Figure 4a). Furthermore, the overall conformation of the carboxylate binding loop between β-strands 1 and 2 of the PDZ domain also contributes to the affinity switch by sterically hindering internal PBM ligands. We therefore considered whether Cdc42-GTP could act as a trigger to release Par6^PDZ^ within the aPKC-Par6-Lgl complex, enabling an apical PBM partner such as Crumbs to bind Par6 (Figure 5a). Consistent with this, a study by Dong et al. indeed proposed that Crumbs was required to switch Par6 from an inhibitory role to an activating role ^11^.

**Figure 5:**
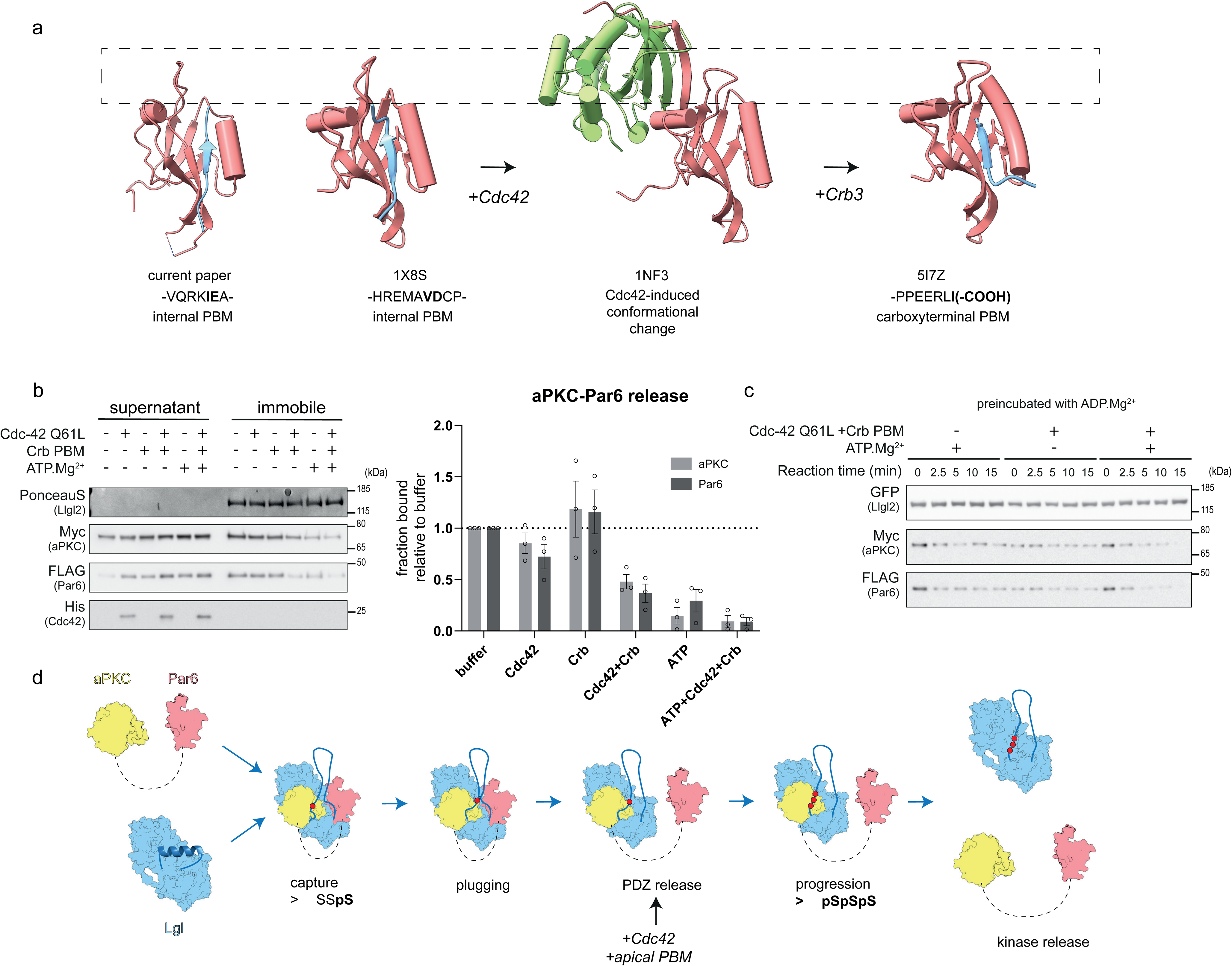
Mechanism of tripartite complex dissolution. (a) Conformational differences between PDZ domains bound to carboxy-terminal or internal PBM ligands, and the conformational change triggered by GTP-bound Cdc42 binding. PDZ domains are coloured in red, PBM ligands in blue and Cdc42 in green. Dashed box is shown to emphasise the carboxylate-binding loop conformation (b) Complex dissolution *in vitro* monitored by Western blot in the presence of the indicated factors. Quantification of the Western blot is shown on the right (n=3 biological replicates represented as mean +/- SEM). (c) Time course of complex disassembly when pre-incubated with ADP-Mg^2+^. Kinetics are shown in Extended data Figure 4b,c (n=4 biological replicates represented as mean +/- SEM). (d) Model for a full phosphorylation cycle for Lgl driven by aPKC-Par6 integrating findings reported here. Lgl is captured through the DPYSD motif interaction with the aPKC kinase domain RIPR motif. The complex is stabilized through the Par6^PDZ^-Lgl^PBM^ contact and an initial phosphorylation event at the C-terminal Ser residue setting up the formation of the plug domain. Further phosphorylation in this stable complex is proposed to be inefficient as the P-site is rotated away from the active site and stabilized by the molecular plug. When the PDZ contact is released by Cdc42-GTP and by a competing PBM-containing protein, the Lgl P-site can be efficiently phosphorylated leading to kinase release (for details see text).

To test these ideas, we co-expressed aPKCι-Par6α together with GFP-Llgl2 in cells, immobilised the tripartite complex on GFP-Trap beads, and then added recombinant Cdc42-GTPγS with a Q61L mutant preventing GTP hydrolysis and/or Crb3 PBM peptide. Comparing the residual bound protein fraction, we observed that the stability of the aPKCι-Par6α-Llgl2 complex was compromised by Cdc42-GTP but also required Crb3 PBM, leading to a 50% reduction in aPKCι-Par6α binding after 20 minutes (Figure 5b). ATP alone also disassembled the complex, presumably by driving phosphorylation to completion (Figure 5b, 2g). Adding both Cdc42/Crb3 and ATP led to an even more efficient release of aPKCι-Par6α with very low residual binding remaining after 20 minutes. These data indicate that Cdc42 and Crb3 PBM can destabilize the tripartite complex and that phosphorylation drives full dissociation. We therefore wondered if Cdc42-mediated PBM release impacted the ATP-driven release of aPKCι-Par6α. In order to probe this effect, we followed complex dissociation kinetics with ATP, Cdc42/Crb3 or both at 16°C (Figure 5c). In these assays, the complex was preloaded with ADP to reflect the product state of the complex after the first reaction step. ATP or Cdc42/Crb3 PBM alone dissociates only a small fraction of the complex under these reaction conditions (Figure 5c, Extended data Figure 4b-d). Adding both Cdc42/Crb3 PBM and ATP to the ADP-preloaded complex resulted in a full release of aPKCι-Par6α after 15 minutes. Importantly when the complex was not preloaded with ADP, ATP itself was sufficient to drive a large proportion of aPKCι-Par6α release, which occurred very rapidly after addition – with about 20% of the complex remaining intact after 2.5 minutes (Extended data Figure 4e-g). This indicates that in the apo state of the complex obtained during immunoprecipitation (∼80%), ATP loading, phosphorylation and release are efficient, while in the ADP-bound state (∼20%), reflecting the likely nucleotide-pocket status of the stalled complex within the cell, requires both Cdc42/Crb3 (Extended data Figure 4e-h). The reaction rate is thus limited by ADP release and we conclude that Cdc42/Crb3 PBM binding to the Par6 PDZ enhances this rate limiting nucleotide release step. The ATP-bound conformation in itself is not sufficient to drive release, as AMP-PNP loading does not result in release of aPKC-Par6 (Extended data Figure 4i). Also the cryo-EM structure was prepared with AMP-PNP, indicating that processive phosphorylation is required for release.

## Discussion

### Mechanistic model for Lgl capture by aPKC-Par6, mutual inhibition and release

Using cryo-EM, biochemical, cellular and *in vivo* experiments, we have identified and validated key interaction sites and their functional roles within the tripartite aPKC-Par-6-Lgl complex, leading to an integrated mechanistic model of complex assembly, mutual inhibition and dissolution. Our data reveal how Lgl is initially captured and oriented in a coordinated fashion by aPKC and Par6 (Figure 5d). aPKC docks onto the second beta-propeller of Lgl using a crucial previously-identified RIPR-docking motif and a large interface consisting of an aPKC-specific C-lobe insert ^23^. The complex is held together by the Par6 PDZ domain interaction with the C-terminal part of the (10-11) loop harbouring an internal PBM motif. A single C-terminal P-site phosphorylation event promotes contacts with aPKC^KD^ proximal to the activation loop in the C-lobe, driving formation of a molecular plug that bridges aPKC and Par6. This plug maintains the N-terminal phosphorylation sites oriented away from the substrate cleft and catalytic site. In this complexed and inhibited state, phosphorylation progression is inefficient as the P-site and PBM motifs are tethered – effectively stalling phosphorylation. When the Par6 PDZ domain dissociates to release the Lgl^PBM^, however, an event that can be triggered by a Cdc42-dependent conformational change (assisted by an apical protein with a high affinity C-terminal PBM), Lgl P-site phosphorylation is allowed to progress. This model explains why interface mutations that block aPKCι docking with Lgl prevent phosphorylation and disrupt monolayer organization and polarity *in vivo*, whereas mutations that perturb Par6 PDZ domain capture of the Lgl protein also do not support maintenance of normal apical-basal organization *in vivo*, yet substantially enhance Lgl phosphorylation (Extended data Figure 5).

Several models (reviewed in Ref 1.) have proposed that the aPKCι-Par6 complex antagonises Lgl by displacing it from the apical membrane by Cdc42-induced phosphorylation and dissolution, but how then does Lgl reciprocally antagonise aPKC in the bound state? Our structure suggests that disassembly of the tripartite complex requires both multi-site phosphorylation of the Lgl P-site *<u>and</u>* release of the Par6^PDZ^ from Lgl. When the Par6 PDZ domain is engaged, we find that the progression of P-site multi-site phosphorylation is impeded. Lgl thus effectively antagonises aPKC-Par6 by trapping it in a tethered product state. We propose that controlled release of Par6 under normal circumstances arises through a Cdc42 induced conformational change and/or competition with binding partners of Par6 at the apical membrane, such as Crumbs ^11,36,37^ leading to plug domain dissolution, rapid ADP to ATP exchange and progression of phosphorylation and disassembly of the tripartite complex. An important implication of this model is that if Lgl encounters aPKC at the basolateral membrane, it inhibits its activity and escorts aPKC away. Such a repressed tripartite aPKC-Par6-Llgl complex would therefore be able to “sense” the apical membrane compartment that contains multiple (competing) partners of Par6 and could respond accordingly unleashing aPKC-Par6 catalytic activity. Our model can also explain the apparently conflicting roles of Par6 as an auxiliary subunit of aPKC – where it is able both to activate and to repress aPKC activity ^11,12^, in addition to contributing to substrate targeting of Lgl.

The precedent set by the existence of the stalled intermediate state that we describe here, as well as the requirement for additional regulatory inputs to facilitate nucleotide exchange has broader implications for kinases. There are numerous examples of ‘processive’ phosphorylation events in nature (reviewed ^38^), and it will be of interest to understand whether some of these are also subject to intermediate product stalled states that are also subject to regulatory input.

In summary, we conclude that the tripartite aPKCι-Par6α-Llgl1 complex that we have characterised structurally and functionally reflects the first cycle or step of the kinase reaction, with Llgl1 monophosphorylated and a hitherto unknown internal Llgl1^PBM^-Par6^PDZ^ interaction that precludes further phosphorylation – stalling processivity. Mutational interrogation of the interfaces that define this complex confirms their critical role in determining polarity. Completion of the reaction cycle based on mutagenesis and reconstitution experiments demonstrate that the subsequent engagement of Cdc42.GTP, switching the Par6 PDZ domain to engage with a Crumbs C-terminal PBM, promotes nucleotide exchange, creating an efficient ATP-dependent completion of phosphorylation and dissociation of the complex. These unprecedented properties revealed through structural analysis provide a comprehensive view of mutual inhibition of aPKC-Par6 and Lgl, the regulatory inputs that determine the dynamics of their engagement and set a series of important precedents informing on kinase-substrate relationships in this system.

## Acknowledgements

We thank members of the McDonald laboratory for helpful comments. We thank Raffaella Carzaniga and Lucy Collinson for EM training. We thank Professor Mark Lemmon and Nate Goehring for critical reading. We thank Natalya Lukoyanova at Birkbeck College for their assistance with data collection. Cryo-EM data were collected at the Institute of Structural and Molecular Biology (ISMB), Birkbeck on equipment funded by the Wellcome Trust, UK (079605/Z/06/Z), and the Biotechnology and Biological Sciences Research Council, UK (BB/L014211/1). This work was supported by the Francis Crick Institute which receives its core funding from Cancer Research UK (CC2068), the UK Medical Research Council (CC2068), and the Wellcome Trust (CC2068). Research in the E.M. lab is supported by FCT—Fundação para a Ciência e a Tecnologia, I.P., (PTDC/BIA-CEL/1511/2021). For the purpose of Open Access, the author has applied a CC BY public copyright licence to any Author Accepted Manuscript version arising from this submission.

## Author contributions

C.P.E. prepared and purified recombinant aPKCι-Par6α-Llgl1 complex, carried out cryo-EM data collection and processing. C.P.E. and N.Q.M. performed the aPKCι-Par6α-Llgl1 complex structure determination and analysis. D.C.B. assisted with the structure refinement. M.C. performed the functional characterization of the complex and the *in vitro* and cell-based experiments. M.C., N.Q.M. and P.J.P. conceived and developed the reaction model. M.E.I. grew crystals of the aPKCι core kinase bound to Llgl2 loop peptide, determined the structure and carried out structure refinement. A.B.C. and E.M. performed all the *in vivo* fly experiments and prepared the related figures. N.Q.M., M.C. and C.P.E. wrote the manuscript. N.Q.M., M.C., E.M. and P.J.P. edited the manuscript.

## Declaration of Interests

The authors declare no competing financial interests.

**Table 1.**
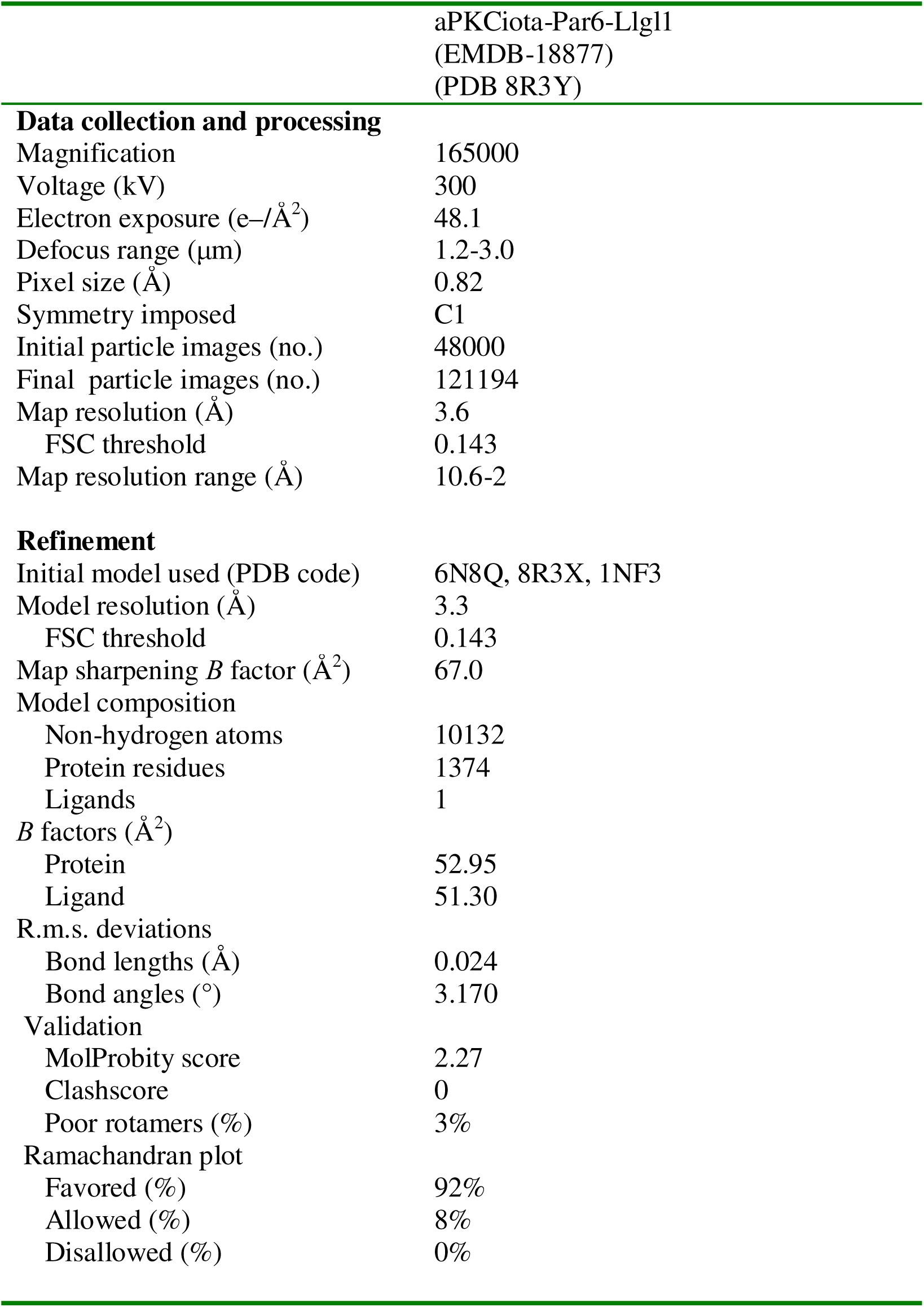
Cryo-EM data collection, refinement and validation statistics.

## Methods

### Cell lines and reagents

HEK293T cells were grown in Dulbecco’s modified eagle medium (DMEM) supplemented with 10% (v/v) Fetal Bovine serum (ThermoFisher Scientific, Waltham, MA, USA), 100U/ml Penicillin and 100µg/ml Streptomycin (ThermoFisher scientific). FreeStyle 293-F cells were grown in Freestyle 293 Expression Medium (ThermoFisher Scientific). DLD1-FlpIn-TREx cells were grown in DMEM supplemented with 10% (v/v) Fetal Bovine serum (ThermoFisher scientific), 100U/ml Penicillin and 100µg/ml Streptomycin (ThermoFisher scientific). DLD1-FlpIn-TREx cells stably harbouring genes for WT and mutant forms of GFP-Llgl2 were selected and grown in the same medium added with 500µg/ml Hygromycin B. Transcription of the transgenes was induced with 200ng/ml doxycycline for 16h unless stated otherwise. Unless stated otherwise, all cloning enzymes were purchased from New England Biolabs (NEB) and other chemicals were purchased from Sigma-Aldrich. GFP-trap agarose was from Chromotek (Planegg, Germany) Anti-Myc (9B11), anti-GFP (4B10), anti-His (Rabbit), phospho-Llgl1/2 S663, secondary HRP-linked goat anti-Rabbit and Horse anti-Mouse antibodies were from Cell Signalling Technologies (Beverly, MA, USA). Anti-FLAG M2 antibody was from Sigma, Phospho-LLGL1/2 S650/654 antibody was from Abgent (San Diego, CA, USA), anti-LLGL1 mAb was from Abnova (Taipei, JP), anti-LLGL2 antibody was from Abcam (Cambridge, UK), Par6B (B-10) antibody was from Santa Cruz (Dallas, TX, USA) Anti-TJP1 antibody was from Atlas (Stockholm, Sweden). Anti-TGN46 antibody was from Abcam (Cambridge, UK) Polyethyleneimine (PEI) was from Polysciences Inc. (Warrington, PA, USA). Mutagenesis and cloning were done using In-Fusion (Takara, Shiga, JP) or Gibson Assembly (NEB). Plasmids and primers used in this study are listed in Table S1. Peptides used were made in-house.

### Protein expression and purification

pCDNA3.1+ plasmids were modified to contain N-terminal Tobacco Etch Virus (TEV) protease-cleavable Twin-Strep-tag or hexahistidine tags. Genes for full-length aPKCι, Par6α, and Llgl1 were amplified by PCR from human cDNAs and inserted into the modified plasmids via Gibson assembly to produce Twin-Strep-tagged Llgl1 and 6xHis-tagged Par6α. Expression plasmids were transfected into FreeStyle 293-F cells (Thermo Fisher Scientific) using linear polyethylenimine (MW 25 000, Polysciences). Cells were harvested by centrifugation after 5 days of shaking at 120 RPM, 8% CO_2_, 37° C. Cell pellets were resuspended in buffer A (20 mM Hepes pH 7.5, 150 mM NaCl, 0.5 mM Tris(2-carboxyethyl)phosphine hydrochloride (TCEP)) supplemented with cOmplete™ EDTA-free Protease Inhibitor Cocktail Tablets (Roche) and lysed by sonication. Cell lysates were clarified by centrifugation and incubated with StrepTactin XT Sepharose (GE Healthcare) for 90 minutes at 4° C. After extensive washing, bound proteins were eluted in buffer A supplemented with 2.5 mM d-Desthiobiotin. Streptactin XT Sepharose eluates were incubated with Ni-NTA agarose (Qiagen) for 1 hour at 4° C in buffer A with 20 mM imidazole prior to extensive washing in the same buffer and elution in buffer A with 250 mM imidazole. Depending on the downstream applications for the sample, tags were either left intact or cleaved off by overnight incubation at 4° C with TEV protease (made in-house). Finally, samples were applied to a Superdex 200 Increase 10/300 GL column (GE Healthcare) in buffer A. Purified proteins were flash-frozen and stored at -80° C.

For expression of the aPKCι kinase domain (aPKCι KD), a recombinant baculovirus was generated for co-expression with phosphoinositide-dependent kinase 1 (PDK1), a priming kinase for aPKCι. Sequences for aPKCι residues 248-596 including an N-terminal GST tag with 3C protease cleavage site and untagged PDK1 were amplified from previous constructs of ours and inserted into the MultiBac pFL vector aPKCι (www.addgene.org). Baculoviruses generated in Sf21 cells using standard protocols were used to infect Sf21 cells at a multiplicity of infection of 2. Cells were harvested by centrifugation after 3 days of shaking at 125 RPM at 27° C and fully-primed aPKCι-KD was purified as described previously ^5^.

### aPKCι-Par6α-Llgl1 cryo-EM grid preparation and data collection

4 µl of aPKCι-Par6α-Llgl1 complex at a concentration of 0.4 mg/ml was incubated with AMP-PNP and applied to R1.2/1.3 Quantifoil 300 mesh copper grids which had been glow-discharged for 45 s at 45 mA. Grids were blotted for 2.5 s at 100% humidity using an FEI Vitrobot MK IV. Data were collected on a Titan Krios transmission electron microscope operated at 300 keV. Data were collected using a Gatan K2 summit direct electron detector operating in counting mode with a GIF Quantum energy filter operating in zero-loss mode. Movies were collected with 8 s exposures dose-fractionated into 40 frames with a total dose of 48.1 e/Å2 and a calibrated pixel size of 0.82 Å. 3407 movies were collected with a defocus range of -3.0 µM to -1.2 µM.

### aPKCι-Par6α-Llgl1 cryo-EM image processing

MotionCor2 and ctffind 4.1 were used for motion correction and CTF estimation, respectively ^39,40^. A total of 3,407 micrographs were selected for further processing. Semi-automated picking with Xmipp3 and particle extraction in Relion-3 yielded 47,516 particles from 1000 micrographs ^41^. After reference-free 2D classification in Relion-3, eight 2D classes were selected and used as templates for reference-based particle picking in Gautomatch [K. Zhang, MRC LMB (www.mrc-lmb.cam.ac.uk/kzhang/)]. A total of 1,069,057 particles were extracted with 2-fold binning and submitted to 8 rounds of 2D classification in Relion-3. After 1 round of 2D classification in CryoSPARC-2, a subset of 48,000 particles was used for *ab initio* 3D model generation in CryoSPARC-2 ^42^. Three models were selected and used as references for 3D classification in Relion-3, this approach yielded 121,194 particles in a single stable 3D class. Particles were re-extracted with the original unbinned pixel size of 0.82 Å in a 280 × 280 pixel box before 3D autorefinement and Bayesian polishing in Relion-3. The polished particles were refined to 3.67 Å resolution using non-uniform refinement in CryoSPARC-2. Finally, the half-maps from CryoSPARC-2 refinement were used as inputs for density modification using Phenix.resolve-cryoem ^43^.

### aPKCι-Par6α-Llgl1 model building

Homology models of each individual Llgl1 β-propeller and the human Par6α PDZ domain were generated using Modeller ^44^ using existing crystal structures of Llgl2 (PDB 6N8Q) and the mouse Par6 PDZ domain (PDB 1NF3), respectively ^13,45^ These models, along with the aPKCι kinase domain crystal structure reported here, were rigid-body docked into the cryo-EM density using UCSF Chimera ^46^. The resulting composite model was subjected to real-space refinement in Phenix using the input models as reference model restraints before manual rebuilding in Coot ^43,47^. The Llgl1 10-11 loop and linkers between the Llgl1 β-propellers were built *de novo* in Coot. Further manual rebuilding in Coot and real space refinement in Phenix yielded a model comprising residues 15-951 for Llgl1, residues 154-252 for Par6α, and residues 248-585 for aPKCι.

### aPKCι-Llgl2 substrate peptide crystallisation and structure solution

aPKCι^KD^ was concentrated to 4 mg/ml and incubated with 1 mM MgCl_2_ and a 3-fold molar excess of both Adenosine 5’-[beta,gamma-methylene]triphosphate (AMP-PCP) and a peptide including residues 644-672 of Llgl2. Crystals grew at 27° C in 25% (v/v) MPD, 25% (v/v) PEG 1000, 25% (v/v) PEG 3350, 0.3 M NaNO_3_, 0.3 M Na_2_HPO_4_, 0.3 M (NH_4_)_2_SO_4_, 0.1 M MES/imidazole pH 6.5. Native data were collected on beamline I04 at Diamond Light Source. Data were scaled using DIALS ^48^, phases were estimated by molecular replacement using Phaser ^49^ with PDB entry 3A8W as a search model. The crystals have two copies of aPKCι-Llgl2 peptide in the asymmetric unit. The Llgl2 substrate peptide was manually built using Coot and the structure was refined at 3.15Å to an R_work_ of 0.218 and an R_free_ of 0.287 with tight geometry using Coot and Phenix ^43,47^. The final model includes Llgl2 residues 657 to 666, AMP-PNP, human aPKCι kinase domain residues 240 to 578.

### Immunoprecipitation and pull-down assays

HEK293T or DLD1-FlpIn-TREx cells expressing GFP-tagged Llgl2 WT and mutants were lysed in 50 mM Tris, pH 7.4, 150 mM NaCl, 1% Triton, 0.5mM TCEP supplemented with phosphatase inhibitors (PhosStop, Roche, Germany), and protease inhibitors (cOmplete, Roche, Germany) and incubated with GFP-Trap magnetic agarose (Chromotek) for 2h at 4°C. Beads were washed once in lysis buffer containing 260mM NaCl and twice in TBS. proteins were eluted in 1xSDS NuPAGE loading buffer (ThermoFisher Scientific). Electrophoresis and western blotting (wet transfer) were done according to standard protocols and imaging was done using a LAS-4000 CCD camera (GE healthcare).

### *In vitro* dissociation assays

To monitor dissociation of Myc-aPKCι and FLAG-Par6α from GFP-Llgl2, the three proteins were co-expressed in FreeStyle293-F cells and GFP-trap was performed as described in the previous section using GFP-Trap magnetic agarose (Chromotek). The immunoprecipitated complex was then added to reaction buffer (20mM Tris, pH7.4, 150mM NaCl, 10mM MgCl_2_,) added with one or more of the following components: Cdc42 (in-house) (1.4μM), GTPγS (100μM), Crb3 PBM peptide (Biotin-Ahx-LPPEERLI-COOH) (100μM), ADP (10μM), ATP (100μM). Reactions were performed at 30°C for endpoint measurements and at 16°C for kinetic measurements. The supernatant was separated from the immobile fraction using a magnet, and the magnetic beads were washed 1x with TBS added with 0.1% Tween-20 before the addition of 2x NuPAGE loading buffer (ThermoFisher Scientific).

### Immunofluorescence staining and confocal microscopy

DLD1-FlpIn-TREx cells stably harbouring genes for GFP-tagged WT and mutant forms of Llgl-2 were seeded on 13_mm glass coverslips at a density of 0.1x10^6^ cells and 24h post plating cultures were induced with 200ng/ml doxycycline for 16h or left uninduced. Cells were fixed with 4% PFA and permeabilized with PBS+0.1% Triton X-100. Coverslips were then blocked in 3% BSA in PBS and incubated with TJP-1 antibody (1:500) or TGN46 antibody (1:500), washed 3x in PBS and subsequently for 2h with goat anti-rabbit 555 (1:1500) or donkey anti-sheep 647 (1:1500) respectively (ThermoFisher Scientific). Coverslips were mounted using Prolong gold with DAPI (ThermoFisher Scientific) and imaged. All the images were acquired using an inverted laser scanning confocal microscope (Carl Zeiss LSM 880 operated using Zen Black software) using a 63x or 40x Plan-APOCHROMAT DIC oil-immersion objective. Images shown in figures were processed in ZEN Blue edition (Zeiss). All images were batch-processed to adjust brightness/contrast. Scoring of ZO-1 staining was done manually by counting cells that were fully enclosed by ZO-1 staining (intact ZO1), displayed partial or fragmented enclosure (discontinuous ZO1) or lacked ZO1 enclosure (loss of ZO1).

### *Drosophila* stocks and genetics

*Drosophila melanogaster* flies were grown using a cornmeal/agar/molasses/yeast media in incubators at temperatures of 18°C and 25°C with controlled photoperiod and humidity. The *lgl^27S3^* null mutant allele (Bloomington *Drosophila* Stock Center (BDSC) #41561) and the following UAS transgenic lines were used: UAS-Lgl-GFP (^32^), UAS-Lgl^ASA^-GFP (this paper); UAS-Lgl^RPYSR^-GFP (this paper); UAS-Lgl^NE^-GFP (this paper). *GR1-GAL4* was used to induce expression of UAS transgenes in the follicular epithelium. The GR1-GAL4 driver shows mild expression during during stages 4 – 7 of oogenesis. All experiments were carried out at 25°C to promote mild-expression mediated by *GR1-GAL4*. The FLP/FRT-mediated mitotic recombination system was used for clonal analyses in the follicle epithelium. Mosaic clones were induced by heat shock at 37°C in flies with the following genotypes:

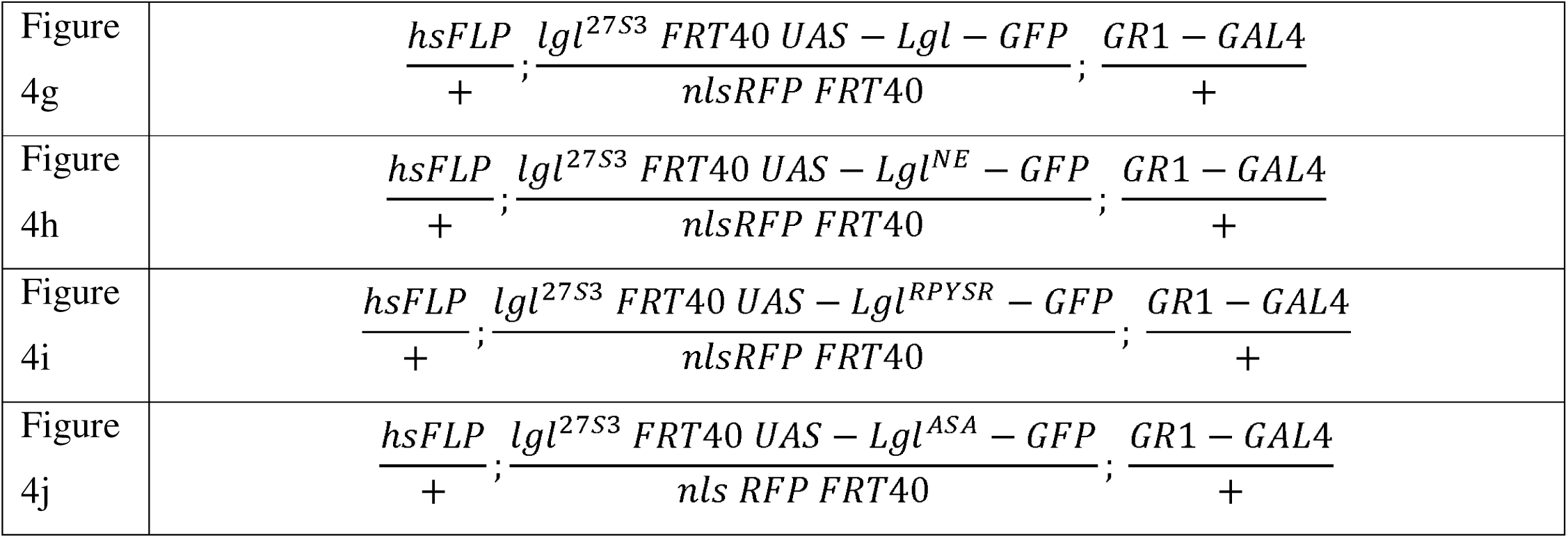

### Cloning and transgenesis of Lgl mutants

Site-directed mutagenesis was performed by Champalimaud Foundation’s Molecular and Transgenic Tools Platform (MTTP) to mutate 502D-502R and 506D-506R (<u>GAT</u>CCTTATTCA<u>GAT</u> to <u>CGT</u>CCTTATTCA<u>CGT</u>) in Lgl^RPYSR^; 695I-695N (<u>ATA</u> to AAC) in Lgl^NE^, and residues 651L-656A and 653R-653A (<u>CTG</u>TCT<u>CGT</u> to <u>GCG</u>TCT<u>GCT</u>) in Lgl^ASA^ using pENTR-Lgl as template ^25^. The GFP-tagged constructs were obtained using LR clonase II to mediate the recombination into pUASt.attb.WG, and were then inserted into the attP-VK18 landing site on chromosome II (BDSC: #9736) via PhiC31 site-specific transgenesis (BestGene Inc). This enables comparable expression levels of the different GFP-tagged mutant versions as all were inserted in the same genomic loci that was used for the control version.

### Fixation and immunofluorescence of *Drosophila* egg chambers

Ovaries of well-fed *Drosophila* females were fixed in 4% paraformaldehyde (in PBS) for 20 min, washed 3×10 min in PBT (PBS with 0.05% of Tween 20), blocked with PBT-10 (PBT supplemented with 10% BSA) and then incubated overnight with primary antibodies in PBT-1 (supplemented with 1% BSA). After 4×30 min washes in PBT-1, ovaries were incubated with secondary antibodies in PBT-0.1 (supplemented with 0.1% BSA) for 2h30, washed three times for 10 min with PBT and mounted in Vectashield with DAPI (Vector Laboratories). The rabbit anti-aPKCzeta (1:500, c-20, Santa Cruz Biotechnology) was used as primary antibody.

### Image Acquisition and analysis in the *Drosophila* follicular epithelium

Immunostainings were analysed using a confocal microscope Leica TCS SP8 (Leica Microsystems) with a PL APO 63x/1.30 Glycerol objective and the LAS X software. Image processing and quantifications were performed using FIJI (Schindelin et al., 2012). Data processing was done in Excel while statistical analysis and graphical representations were performed using GraphPad Prism 8 tools. To analyse the ability of different Lgl mutants to recapitulate Lgl function on epithelial apical-basal organization, we monitored if expression of Lgl transgenes carrying different mutations would rescue the fully penetrant multilayering phenotype of tissue homozygous for the *lgl^27S3^* null allele ^32^. Multilayering (defined has 3 or more epithelial cells piling on top of the epithelial layer) was scored by inspecting midsagittal cross-sections of stage 4 to stage 7 egg chambers. Only egg chambers with large mutant clones (> 1/4 of the whole egg chamber mutant) were considered in the analysis. The developmental stage of egg chambers was determined by measuring their area in midsagittal cross-sections, as a proxy for size. To define the area intervals corresponding to each developmental stage, we staged control egg chambers stained for aPKC and overexpressing UAS-Lgl::GFP flies according to phenotypic characteristics and correlated area size with the developmental stage. For each independent experiment the analysed egg chambers were obtained from a minimum of 10 flies per genotype.

### Data availability

The cryo-EM map of aPKCι-Par6α-Llgl1 complex is available in the Electron Microscopy Data Bank (accession number EMD-18877). The structure coordinate file for the fitted aPKCι-Par6α-Llgl1 model is available in the Protein Data Bank database (accession number 8R3Y). The structure coordinate file for the fitted aPKCι kinase domain bound to Llgl2 P-site peptide is available in the PDB database (accession number 8R3X). All biological materials generated in this manuscript are available from the authors upon request. Further information on the research design is available in the Nature Research Reporting Summary linked to this article. Source data are provided with this paper.

