## Extended data figure legends for "Capture, mutual inhibition and release mechanism for aPKC-Par6 and its multi-site polarity substrate Lgl"

**Extended Data Figure 1. Purification and cryo-EM structure determination of an aPKCι-Par6α-Llgl1 complex**

(a) Schematic of aPKCι-Par6α-Llgl1 complex and location of affinity purification tags. Representative SEC profile for aPKCι-Par6α-Llgl1 monitored by SDS-PAGE across gel filtration fractions. (b) Cryo-EM processing workflow for aPKCι-Par6α-Llgl1 complex. Data collection statistics are shown. 0.82Å/pixel size was used and data were collected using a K2 detector on a Titan Krios microscope. A motion-corrected cryo-EM micrograph of the aPKCι-Par6α-Llgl1 complex is shown. The processing workflow shows a representative selection of cryo-EM 2D class-averages of aPKCι-Par6α-Llgl1 calculated using Cryosparc after using final 3D classification. Three 3D models were generated for which 121,000 particles contributed to the higher resolution 3D reconstruction. The structure was determined to an average resolution of 3.6 Å as judged by a Fourier-shell correlation (FSC=0.143) criterion as shown. (c) Local resolution estimation. Shown are a local filtered map from CryoSparc2 (pale blue) and a density modified and auto-sharpened map from PHENIX indicating local resolution. (d) angular distribution of particles used in final reconstruction (e) Top view of model fitted to resolve-sharpened map contoured at 0.7 (f) Back view of model fitted to resolve-sharpened map contoured at 0.7 (g) Cryo-EM map density (dark blue chicken wire) superposed with final model within the aPKCι nucleotide pocket indicating bound AMP-PNP (h) Location of the three phospho-sites aPKCι ^pT412^, aPKCι^pT654^ and Llgl1^pS663^. Density for each modification superposed with final model for (i) the aPKCι turn motif pT654 residue, (j) aPKCι^pT412^ within the the aPKCι activation loop and (k) the Llgl1 pS663 residue within the P-site. (l) Regions not observed in the final structure (PB1 domains from aPKCι and Par6α, the pseudo-substrate and C1 domain of aPKCι) are greyed out, while poorer density proximal to the aPKCι N-lobe is not interpreted (dark grey density).

**Extended Data Figure 2. Multisite phosphorylation of Lgl**

(a) Binding affinity of the Llgl1 P-site peptide measured by fluorescence anisotropy. (b) Crystal structure of the Llgl2 substrate peptide with the C-terminal Ser residue positioned as the phospho-acceptor, with residues numbered from Ser at residue 0. Superposed is the final electron density map calculated using sigma-A weighted mFo-DFc coefficients. The ordered part of the Llgl2 P-site is shaded orange with the -2R and S phospho-acceptor site indicated by an underline. (c) Specificity of phospho-Llgl antibodies used in this study. Western blot analysis using phospho-specific antibodies against pS653 site or pS645/pS649 sites for different Llgl2 phospho-acceptor site serine mutations in the P-site of loop (10-11). Phosphorylation of WT and the indicated mutants of Llgl2 is shown upon co-expression with aPKCι. The Western blot shows the specificity of the two different phospho-specific antibodies against either the pS653 site or pS645/pS649 sites (Llgl2). Representative western blot of n=2 experiments.

**Extended Data Figure 3. Properties of Llgl2 mutants in DLD1 cells**

(a) Co-localization of Llgl-2^IE>NE^ in DLD1 cells with TGN-46. A subfraction of doxycycline (Dox)-induced Llgl-2^IE>NE^ protein localizes with foci staining positive for TGN-46. Representative micrographs shown from one of two separate cover slips (b) Localization of Llgl2 RPYSR, LSR>ASA, or SSS>AAA mutants of Lgl-2 in DLD1 cells and the effect of ectopic protein expression on ZO-1 staining as a marker of cell polarity. GFP-tagged WT or mutant forms of Llgl-2 were expressed in DLD1 cells via doxycycline induction. Llgl-2 and ZO-1 localization were monitored by confocal microscopy. Representative micrographs shown for one of three independent biological replicates (c) Quantification of ZO-1 staining patterns observed in (b). (d) Phosphorylation of endogenous Llgl1/2 with and without doxycycline induction. Overexpression of WT-Llgl2 suppresses the phosphorylation of endogenous Llgl1/2 in a dominant fashion, whereas the expression of Llgl-2^IE>NE^ has no effect on the phosphorylation levels of endogenous protein, while it is hyperphosphorylated compared to WT GFP-Llgl2. Representative western blot of n=5 biological replicates.

**Extended Data Figure 4. Release of Lgl from aPKC-Par6α is driven by Cdc42, Crb and ADP-ATP exchange**

(a) Steric clash between K162 (human Par6α numbering) of the Par6 PDZ domain in the Cdc42-induced conformation (5I7Z) shown in purple and P712 from a structural superposition of the Llgl-1 internal PBM (this study) shown in blue. The clash indicates that the Cdc42-induced conformation is incompatible with a bound internal PBM. Cdc42 is coloured in green and the Par6α PDZ domain of the reported structure in red. (b) aPKCι release kinetics quantified from the aPKCι-Par6α-Llgl2 complex Western blots as shown in Figure 5c. Quantified from n=4 biological replicates represented as mean +/- SEM (c) Par6 release kinetics shown in Figure 5c. Quantified from n=4 biological replicates represented as mean +/- SEM. Points fitted to a one phase exponential decay dissociation curve using GraphPad Prism (d) Schematic representation of the predicted proportions of tripartite complex with aPKCι in the apo or ADP-bound state when preloaded with ADP.Mg^2+^ and their respective trajectories for release. Proportions inferred from residually bound protein levels in panels b and c. (e) Time course of aPKCι-Par6 release from Llgl2 and the influence of the indicated factors without preincubation with ADP.Mg^2+^. Representative western blot of n=2 biological replicates. (f) aPKCι release kinetics quantified from Western blots and shown in e. Quantified from n=2 biological replicates represented as mean +/- SEM (g) Par6 release kinetics as in e. Quantified from n=2 biological replicates represented as mean +/- SEM. Points fitted to a one-phase exponential decay dissociation curve using GraphPad Prism (h) Schematic representation of the predicted proportions of complex obtained as apo or ADP-bound and their respective trajectories for release. Proportions inferred from residually bound protein levels in panels f and g. (i) Time course of complex disassembly induced by ATP.Mg^2+^ versus AMPPNP.Mg^2+^ when pre-incubated with ADP-Mg^2+^. Representative western blot of n=3 biological replicates.

**Extended Data Figure 5. Impact of disruptive Llgl1/2 mutations on tripartite complex behaviours**

Explanation of the behaviours and properties of four disruptive Llgl1/2 mutations described in the text, referring to the overall model presented in Figure 5. The IE>NE mutant in the PBM bypasses the plugged state of the complex, leading to rapid phosphorylation progression and complex dissolution. The LSR>ASA mutant of the high-affinity kinase docking motif is less stably tethered to the kinase domain as the wild-type protein, resulting in more rapid release combined with less efficient N-terminal Ser phosphorylation, while C-terminal phosphorylation is unaffected. The RPYSR mutant displays a decreased kinase domain interaction and is unable to correctly position the kinase domain for efficient phospho-transfer, resulting in an overall suppression of phosphorylation. The SSS>AAA non-phosphorylatable mutant is trapped in the capture state as it cannot be phosphorylated to form the initial product state required to assemble the plug. It behaves similar to the WT protein in overexpressed conditions, which is trapped in the plugged state, as it likely saturates the endogenous release mechanism impeding the progression of the reaction.
