## Supplementary figures and images for "Capture, mutual inhibition and release mechanism for aPKC-Par6 and its multi-site polarity substrate Lgl"

### Extended data figure 1

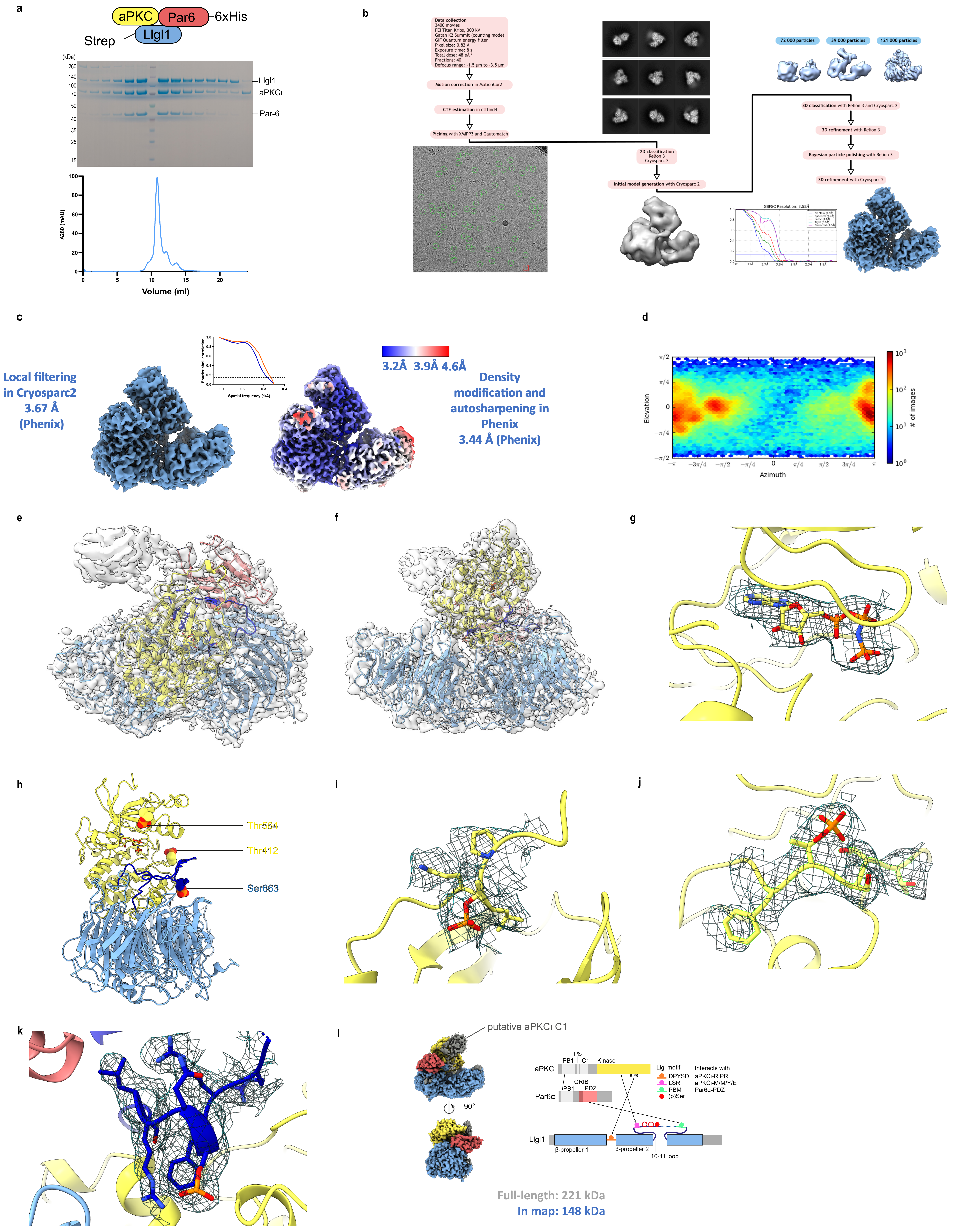

### Extended data figure 2

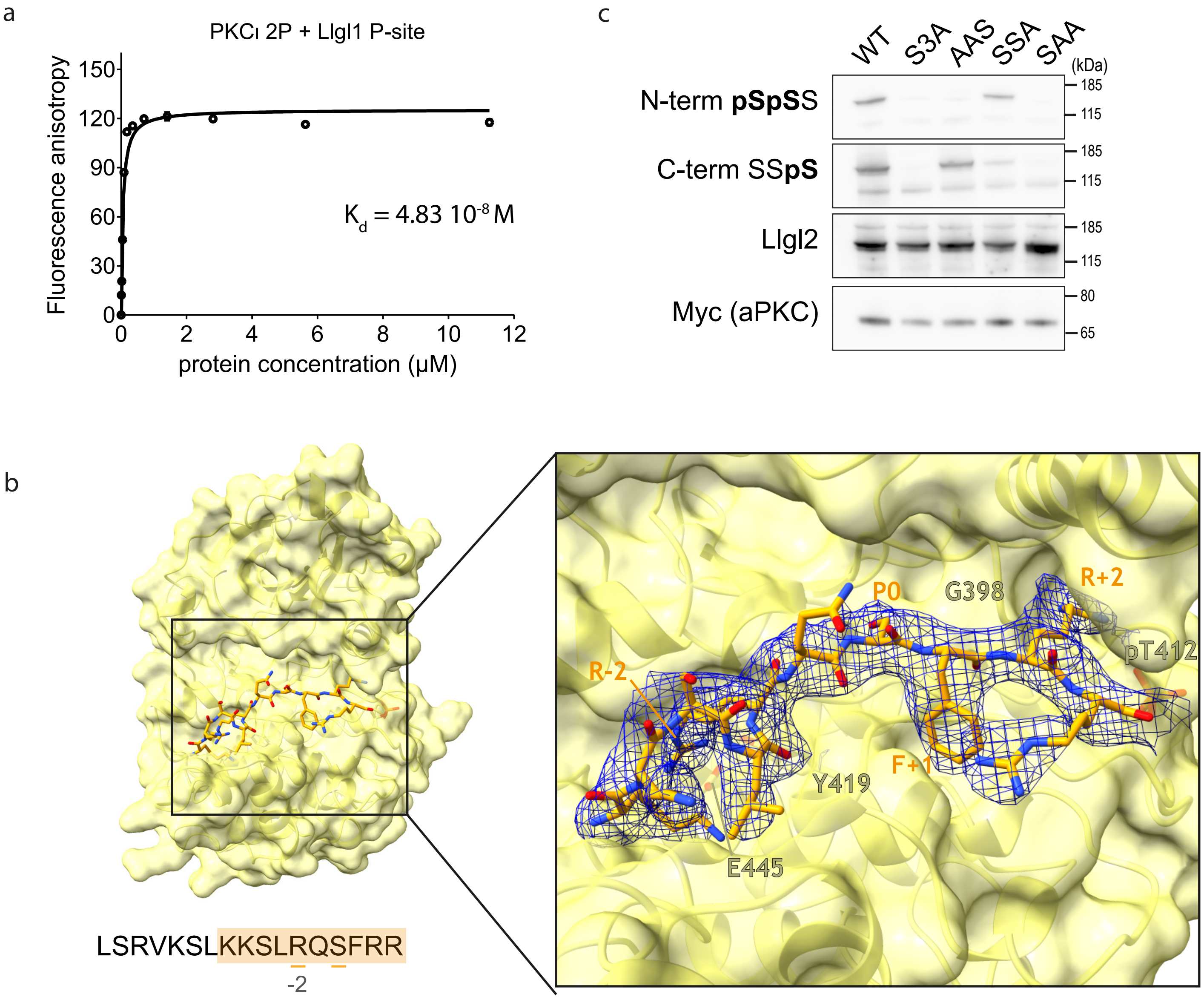

### Extended data figure 3

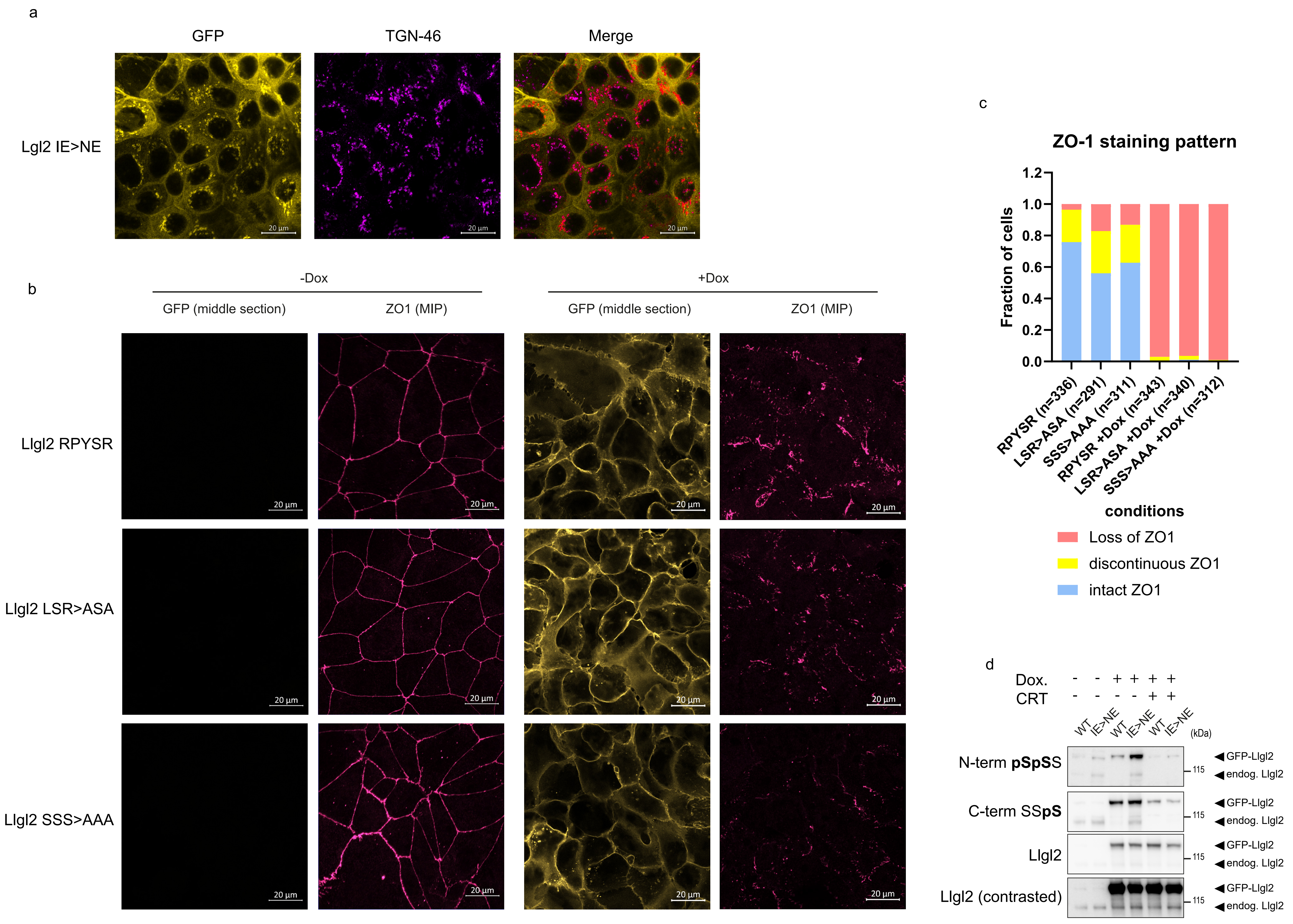

### Extended data figure 4

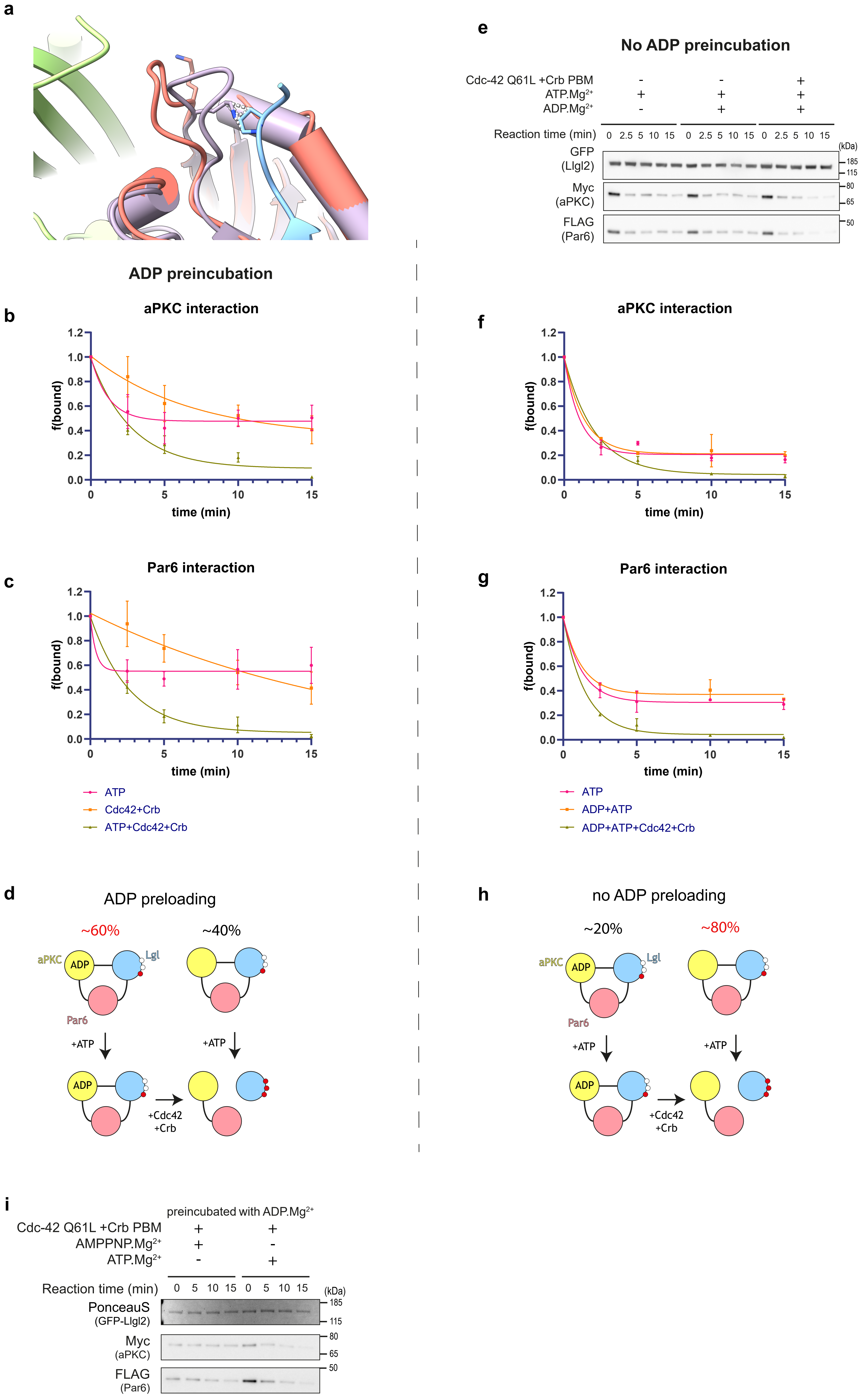

### Extended data figure 5

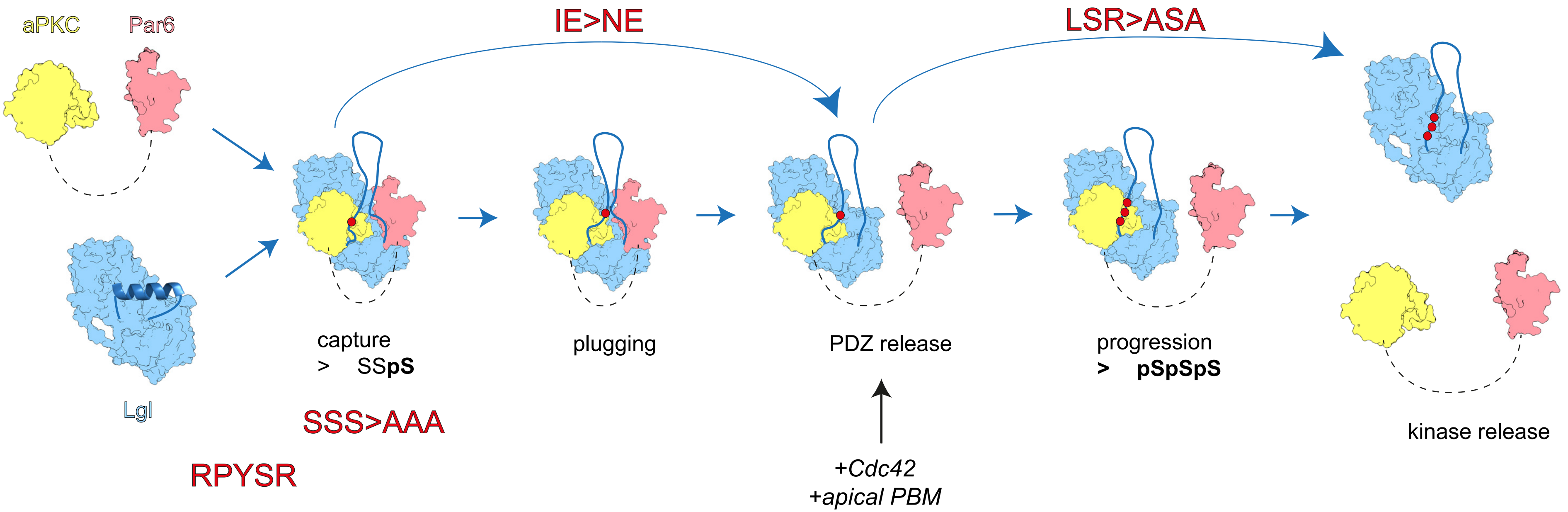
